# TRIOBP promotes bidirectional radial stiffness gradients within the organ of Corti

**DOI:** 10.1101/2021.06.28.450090

**Authors:** Hesam Babahosseini, Inna A. Belyantseva, Rizwan Yousaf, Risa Tona, Shadan E. Hadi, Elizabeth Wilson, Shin-ichiro Kitajiri, Gregory I. Frolenkov, Thomas B. Friedman, Alexander X. Cartagena-Rivera

**Author notes:** Co-first authors (Hesam Babahosseini, Inna A. Belyantseva). To whom correspondence should be addressed at: A.X.C.R., NIBIB/NIH, Bethesda, MD, 20892, USA.

## Abstract

Hearing depends on complex mechanical properties of the inner ear sensory epithelium. Yet, the individual contributions of different cell types to the stiffness spectrum of the sensory epithelium have not been thoroughly investigated. Using sub-100 nanometer spatial resolution PeakForce Tapping Atomic Force Microscopy (PFT-AFM), we mapped the Young’s modulus (stiffness) of the apical surface of different cells of freshly-dissected cochlear epithelium from wild-type mice and mice lacking the F-actin bundling protein TRIOBP-5 or TRIOBP-4 and TRIOBP-5. Variants of the genes encoding human and mouse TRIOBP are associated with deafness. We show that TRIOBP deficiency affects formation of supporting cell apical phalangeal microfilaments and bundled cortical F-actin of hair cell cuticular plates, softening the apical surface of the sensory epithelium. Unexpectedly, high-resolution PFT-AFM-mapping also revealed previously unrecognized reticular lamina radial stiffness gradients of opposite orientations in wild-type supporting and hair cells. Deafness-associated TRIOBP deficiencies significantly modified these bidirectional radial stiffness gradients.

## Introduction

In the inner ear, the sensory epithelium of the organ of Corti detects and amplifies sound vibrations with exquisite sensitivity. The specialized cell types of the organ of Corti have intricate morphologies and together build an ultrasensitive electrochemical and mechanical machine. Mechanotransducing hair cells are flanked by supporting cells providing resistance to mechanical deformations from sound stimulation [1] and yet allow propagation of sound waves. Sensory cells in the cochlea are organized in rows along the cochlear spiral with one row of inner hair cells (IHCs) and three rows of outer hair cells (OHCs) (Fig. 1). The sensory IHCs transduce sound and convey auditory information directly to the brain’s auditory nuclei, while OHCs amplify weak sound-evoked vibrations [2]. Supporting cells in the organ of Corti include inner pillar cells (IPCs), outer pillar cells (OPCs) and three rows of Deiters’ cells (DCs) (Fig. 1). The apical surfaces of IPCs are located between the IHCs and OHCs, while OPCs extend their apical surfaces between the first row of OHCs and make a contact with the second row of OHCs. The apical plates and processes of DCs separate the other two rows of OHCs (Fig. 1). The apical plates of DCs and OPCs are the main constituents of the reticular lamina (Fig. 1) that extends from the heads of OPCs to the Hensen’s cells and immobilizes the OHC cuticular plates. Together with OHCs, these supporting cells are interconnected by tight junctions that provides boundaries between the endolymph and perilymph compartments and prevents mixing of these two fluids that have very different ionic compositions [3]. The reticular lamina also provides structural support and facilitates the exceptional sensitivity of hair cells required for normal sound transduction [4].

**Fig. 1.**
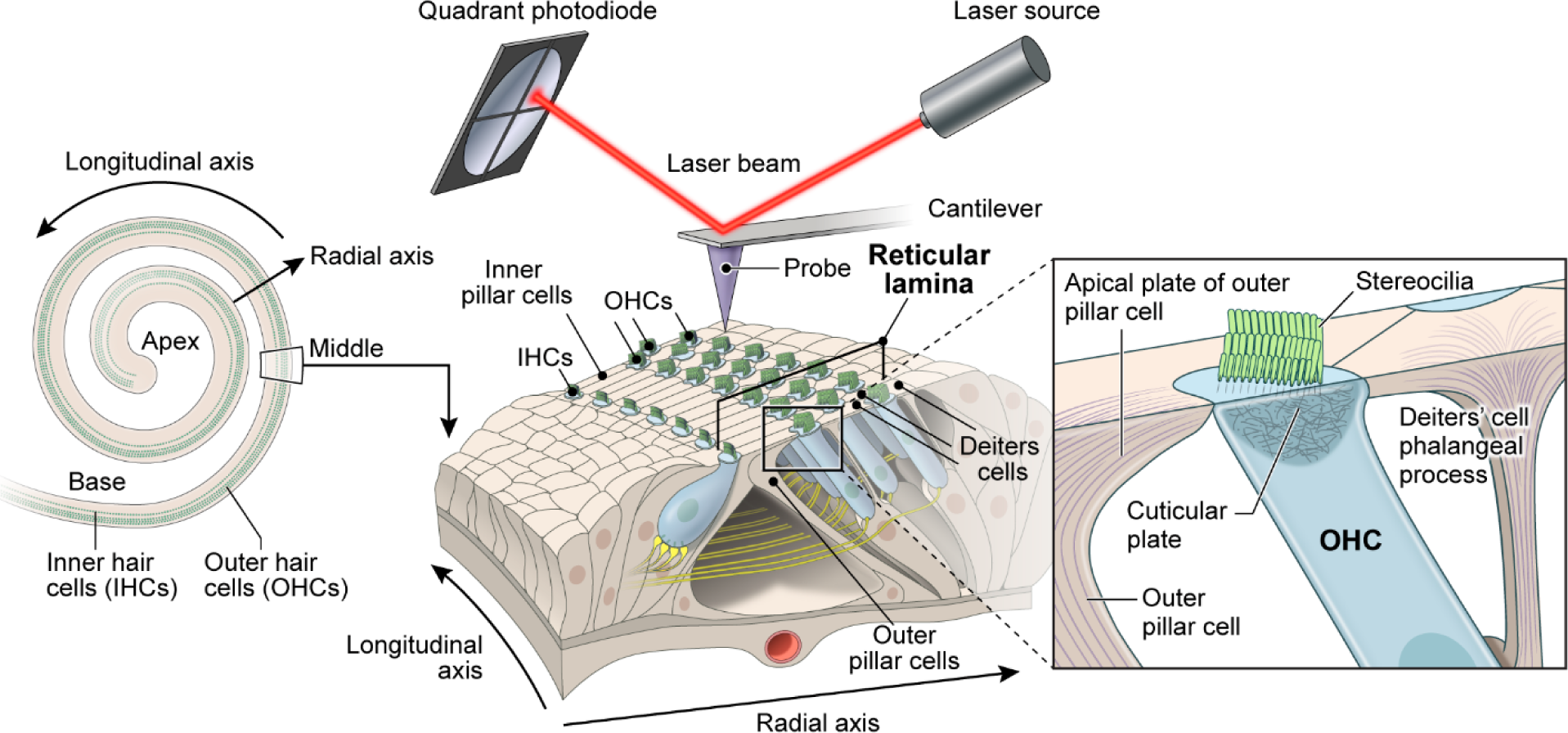
Schematic of the organ of Corti sensory epithelium, hair cells, supporting cells cellular structures, and the experimental system for PFT-AFM measurements. **a** The schematic illustrates the coiled shape of the inner ear auditory sensory epithelium. **b** The PFT- AFM experimental setup that was used to measure nanoscale stiffness of the organ of Corti reticular lamina of sensory epithelium segment including three rows of OHCs and the supporting cells (IPCs, OPCs, and three rows of DCs). **c** Close-up view of the longitudinal cross-section through the OHC and surrounding supporting cells showing the F-actin-based cuticular plate, stereocilia bundle of OHC, OPC apical plate and DC’s phalangeal process.

To transduce sound, each hair cell has a mechano-receptive hair bundle with stereocilia rows of increasing height organized in a staircase arrangement [5]. In response to sound, a stereocilia bundle is deflected a few tens to 100 nanometers, which opens tension-gated mechanoelectrical transduction ion channels located at the tips of shorter row stereocilia [6]. Each stereocilium has a paracrystalline filamentous actin (F-actin) core, consisting of hundreds of unidirectionally oriented, bundled and cross-linked actin filaments [5]. Adjacent stereocilia in a bundle are interconnected by a variety of extracellular links [5]. Each stereocilium is anchored into the apical hair cell body by an F-actin-based rootlet that is embedded in the cuticular plate, a rigid filamentous actin cytoskeletal structure located below the apical plasma membrane of a hair cell [7] (Fig. 1). Stereocilia are rigid structures. When deflected by sound waves, all stereocilia within a hair bundle bend at their insertion points in the hair cell [8]. A rootlet is embedded in the F-actin stereocilium core from approximately one third to one half the stereocilium length and about the same distance in the opposite direction into the cuticular plate where each rootlet is anchored [7].

The elaborate F-actin cytoskeleton of hair cells requires many actin-bundling proteins. One of them that is essential for stereocilia rootlet development and function is TRIOBP, which is also required as an important component of the apical cytoskeleton of adjacent supporting cells [9, 10]. There are three classes of alternative splice mRNA isoforms transcribed from the mouse *Triobp* and human *TRIOBP* genes encoding the proteins TRIOBP-1, TRIOBP-4, and TRIOBP-5 [9]. Mouse *Triobp-4* mRNA is transcribed from the first eight exons of the *Triobp* gene and encodes a 107-kDa protein with unusual repeated motifs that bind and tightly bundle actin filaments [11]. TRIOBP-5 is a 218-kDa protein [11], which includes all of the amino acid sequence of TRIOBP-4 at its N-terminus while the remainder of TRIOBP-5 has five predicted coiled-coil domains and a pleckstrin homology (PH) domain [9, 10, 12]. TRIOBP-1 is the smallest of the three isoforms (72-kDa) and is also named Tara [12]. *Triobp-1* mRNA has a unique first exon with its own transcription start site located between exons 12 and 13 of the *Triobp* gene, while the rest of the *Triobp-1* mRNA is identical to exons 13 through 25 of *Triobp-5* mRNA. TRIOBP-1 protein binds and stabilizes F-actin structures [12-15], shows a broad expression pattern throughout the body and, at least in mouse, is required for embryonic viability. However, TRIOBP-4 and TRIOBP-5 are only necessary for normal hearing [9, 10, 16-21]. Most of the recessive variants of human *TRIOBP* are associated with profound deafness DFNB28 and are located in the large exon 6, orthologous to mouse exon 8, and thus they mutate simultaneously both TRIOBP-4 and TRIOBP-5 (TRIOBP-4/5) proteins [16, 17]. To date, variants that damage only the TRIOBP-5 isoform are associated in human with moderate to severe deafness or age-related hearing loss [18-20].

In mouse inner ear hair cells, TRIOBP is mainly localized to stereocilia rootlets [9, 10]. Deficiency of both TRIOBP-4 and TRIOBP-5 (TRIOBP-4/5) results in a total absence of rootlets. However, a deficiency only of TRIOBP-5 results in dysmorphic stereocilia rootlets and a progressive loss of hearing [9, 10, 22]. TRIOBP is also expressed in supporting cells of the sensory epithelium and is localized in the apical processes of IPCs, OPCs, and three rows of DCs. We recently reported a critical role of the TRIOBP-5 protein in supporting cell mechanics [10]. Exactly how the overall micromechanics of the cochlear reticular lamina depends on the functions of different TRIOBP isoforms is understudied. Understanding the composite of TRIOBP functions in cochlear mechanics requires a comprehensive investigation of the mechanical properties of the different individual cell types in the inner ear at nanoscale resolution. We hypothesize that the highly divergent cell types in the organ of Corti show multidimensional but interconnected mechanical relationships.

The mechanical resistance of individual cells interacting in a live organ of Corti is critical for sound wave propagation and transduction. In previous studies, the cochlear micromechanical properties at the cellular level have been measured in spatially restricted areas, such as the stereocilia bundles, using glass micro-pipettes [23, 24], fluid jets [25] or standard atomic force microscopes [26]. Quasi-static AFM was also utilized to characterize the surface Young’s modulus of OHCs and supporting PCs in the mouse cochlea at early ages (P0-P5) [27]. Due to the low spatial resolution of these studies, the cochlear mechanical properties were not mapped. Changes in Young’s modulus of the cochlear basilar membrane using PeakForce Tapping Atomic Force Microscopy (PFT-AFM) technique was reported [28]. However, to date, no study has systematically measured in a holistic manner at nanoscale resolution the mechanical properties of the spatially heterogenous apical surface of the cochlear sensorial reticular lamina of wild-type and mutant mice with cytoskeletal alterations of the different cells of the organ of Corti.

In this study, we used PFT-AFM to measure the axial elastic modulus of the reticular lamina including individual OHCs at the cuticular plates and stereociliary bundles, the apical surface of the PCs and the apical processes of DCs (Fig. 1). We also investigated how a combined TRIOBP-4 and TRIOBP-5 deficiency, or a TRIOBP-5 deficiency alone affects the apical stiffness of individual cells. In doing so, we identified additional functions of TRIOBP-4 and TRIOBP-5 in the modulation of the reticular lamina micromechanics in the organ of Corti. Intriguingly, we discovered previously unknown radial stiffness gradients of opposite orientations for supporting cells and hair cells. The absence of both TRIOBP isoforms significantly reduces the magnitude and affects the orientation of these radial stiffness gradients, which are essential for optimal tissue-level nanomechanics of the auditory sensory epithelium.

## Results

### Quantification of Triobp-4 and Triobp-5 mRNAs expressed in mouse organ of Corti

TRIOBP has multiple cytoskeletal functions in different inner ear cell types. The expression of mouse TRIOBP-4 and TRIOBP-5 was examined using a LacZ cassette (*E. coli β*-galactosidase) that replaced all of the sequence of mouse *Triobp* exon 8. Therefore, *β*-galactosidase cellular expression is a proxy for both TRIOBP-4 and TRIOBP-5 proteins because exon 8 sequence is included in both *Triobp-4* and *Triobp-5* mRNAs. Staining for *β*-galactosidase activity in the sensory epithelium of the inner ear [9], however, could not discriminate the *Triobp*-driven LacZ signal for *Triobp-5* from a signal of *Triobp-4* expression. Therefore, rather than a LacZ reporter, in this study we examined *in situ Triobp* mRNA expression using RNAscope probes specific for (1) both *Triobp-4 and Triobp-5* mRNAs (*Triobp-4/5*) (Fig. 2a, left and middle panels), (2) both the *Triobp-1* and *Triobp-5* mRNAs (*Triobp-1/5*) (Fig. 2b, left and middle panels) and (3) only the *Triobp-*5 mRNA (Fig. 2a and b, right panels). In wild-type mouse cochlea at P6, *Triobp-5* mRNA expression is prominent in outer hair cells and to a lesser extent in inner hair cells and supporting cells. A similar pattern was observed when a RNAscope probe specific for *Triobp-1/5* mRNA was utilized. In addition to the hair cell signal, there was a weaker signal in supporting cells. In contrast, an RNAscope probe specific for *Triobp-4/5* mRNAs shows a more broadly distributed signal highlighting both hair cells and supporting cells, including cells of the greater epithelial ridge (Fig. 2a, middle panels). This pattern of expression of *Triobp-4/5* mRNA is consistent with the reported role of TRIOBP-4, which is necessary for stereocilia rootlet formation and the initial development of the cytoskeletal components of supporting cells required for their structural role in the reticular lamina [9]. The pattern of TRIOBP-5 mRNA expression at P6 is consistent with the reported conclusion that this isoform plays major roles in the maturation of stereocilia rootlet architecture and in the development of cytoskeletal structures of the supporting cells, which becomes critically important after the onset of hearing at P12 [9, 10].

**Fig. 2.**
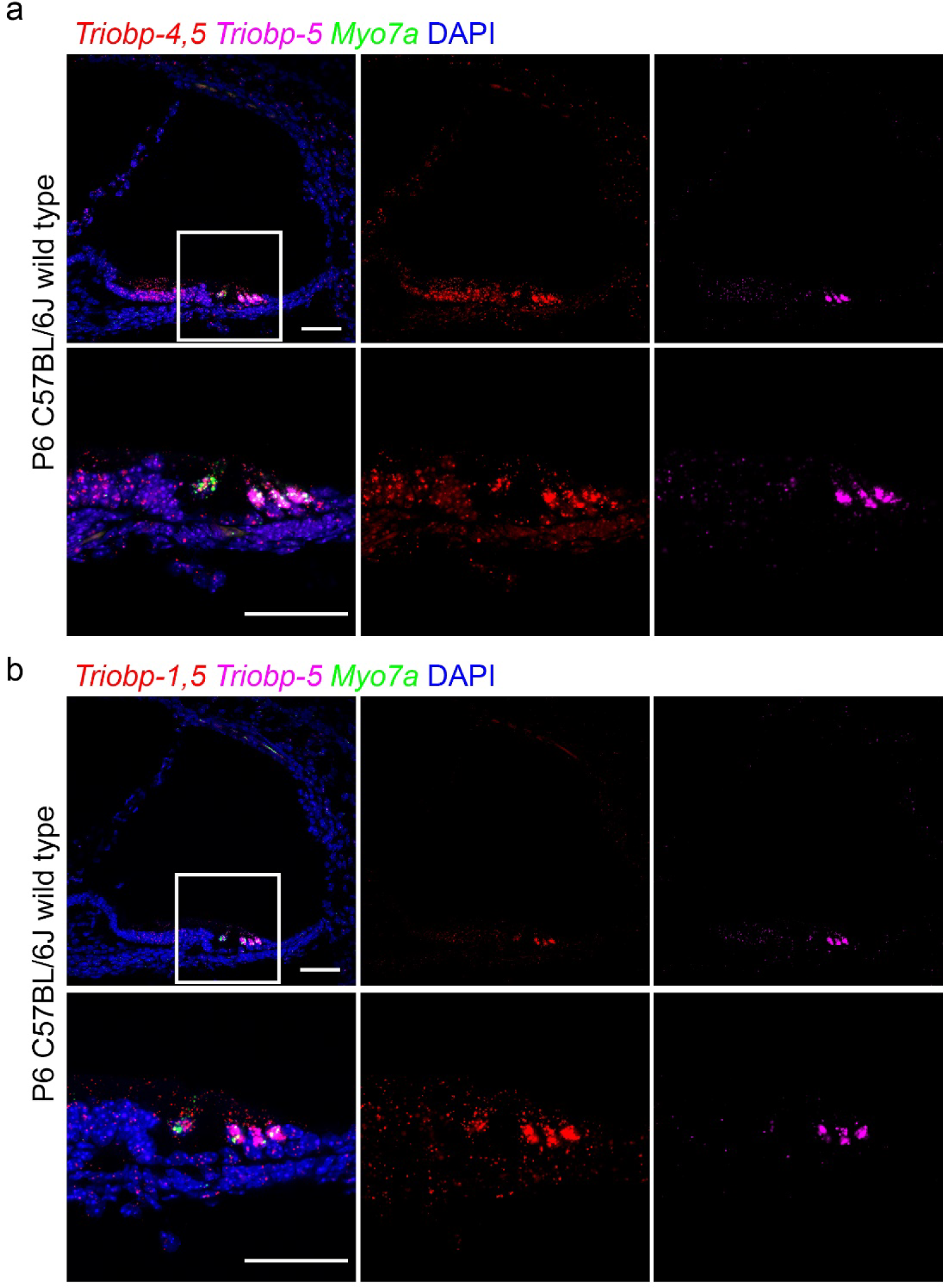
*In situ* hybridization using RNAscope probe in P6 wild-type mouse cochlea. **a** Expression of *Triobp-4 and Triobp-5* mRNAs (*Triobp-4/5*) (red, Probe-Mm-Triobp-O1), *Triobp- 5* only mRNA (magenta, Probe-Mm-Triobp-O2-C3) and *Myo7a* mRNA (green, Probe-Mm- Myo7a-C2). *Triobp-4/5* mRNA (red) is expressed in inner and outer hair cells and supporting cells. Whereas mRNA of *Triobp-5* alone is expressed mainly in OHCs. **b** Expression of both *Triobp-1 and Triobp-5* mRNA (*Triobp-1/5*) (red, Probe-Mm-Triobp-O3), *Triobp-5* only mRNA (magenta, Probe-Mm-Triobp-O2-C3) and *Myo7a* mRNA (green, Probe-Mm-Myo7a-C2). *Triobp- 1/5* mRNA was detected mainly in hair cells. Scale bars are 50µm.

Quantification of expression levels of the three different *Triobp* isoforms in the inner ear of P6 *Triobp^ΔEx9-10^* mutant mice, deficient only for the TRIOBP-5 isoform, was also evaluated using droplet digital PCR (ddPCR). In heterozygous *Triobp^ΔEx9-10/+^* mice, there was a reduction of *Triobp-5* expression level to approximately one-half of the wild-type. These data indicate that there is no increase in mRNA expression from the remaining single wild-type copy of the *Triobp* gene to compensate for the loss of one allele in heterozygotes. As expected, no expression of *Triobp-5* mRNA was detected in the homozygous *Triobp^ΔEx9-10 /ΔEx9-10^* mouse (Supplementary Fig. 1b). In addition, no significant difference in the expression level of either *Triobp-1 or Triobp-4* mRNAs was observed in the homozygous *Triobp^ΔEx9-10/ ΔEx9-10^* mice, indicating that the *Triobp^ΔEx9-10^* allele does not induce an up-regulation or down-regulation of the *Triobp-1* or *Triobp-4* mRNAs (Supplementary Fig. 1b) and by inference their protein levels.

### TRIOBP-5 deficiency results in ultrastructural defects in OHCs, OPCs and DCs

The effects of TRIOBP-deficiency on hair cell stereocilia rootlet formation and architecture are published [9, 10]. However, the ultrastructural changes in the other regions of hair cells and in supporting cells have not been investigated by electron microscopy. To explore these potential changes, we used serial sectioning with Focused Ion Beam and Scanning Electron Microscopy imaging (FIB-SEM) that provides a lateral X-Y resolution of approximately 2nm and a sectioning step of 20nm in the high-pressure frozen freeze-substituted cochlear explants. In heterozygous *Triobp^ΔEx9-10/+^* mice at P6, virtual transverse re-slicing of an OHC in the registered FIB-SEM stack revealed regular filamentous actin “tangles” at the top of the cuticular plate. These regular tangles were not observed in homozygous TRIOBP-5 deficient littermates (Fig. 3a). The disruption of cuticular plate F-actin in *Triobp^ΔEx9-10/ΔEx9-10^* OHCs represents either a direct effect of TRIOBP-5 deficiency or F-actin re-arrangement associated with the abnormalities of stereocilia rootlets in these mice [10].

**Fig. 3.**
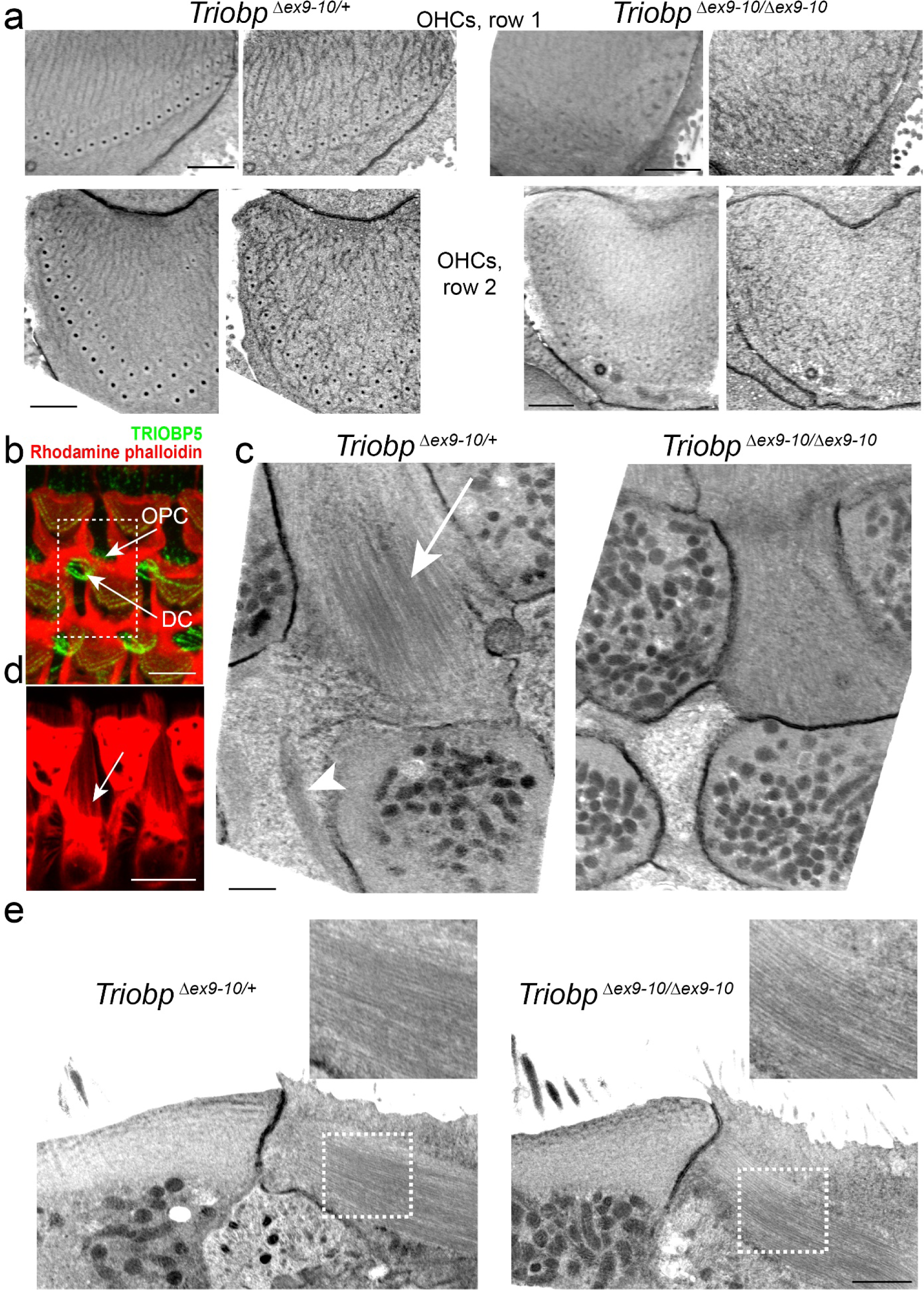
Ultrastructural effects of TRIOBP-5 deficiency. **a** Regular actin “tangles” at the upper surfaces of the OHC cuticular plates in *Triobp^ΔEx9-10/+^* mice (left) and their disruption in *Triobp^ΔEx9- 10/ΔEx9-10^* mice (right). Actin filaments were stabilized with tannic acid during fixation before staining with uranyl acetate in the freeze-substitution step (see Methods). Upper images illustrate OHCs1 (first row OHCs) while bottom images illustrate OHCs2 (second row OHCs). Each cuticular plate is shown first as a median Z-projection of a 400nm-thick FIB-SEM volume and then as a single representative FIB-SEM section. The data are representative of five *Triobp^ΔEx9-10/+^* OHCs and eight *Triobp^ΔEx9-10/ΔEx9-10^* OHCs. **b** Example of TRIOBP-5 immunofluorescent labeling (green) and F-actin rhodamine phalloidin labeling (red) in the outer pillar cells (OPC) and Deiters’ cells (DC). **c** Median Z-projections of a 400nm-thick FIB-SEM volume at the level of the bottom of the OHC cuticular plates, which show microfilaments/microtubules in OPCs (arrow) and actin patches in DCs (arrowhead) in *Triobp^ΔEx9-10/+^* mice (left) and their disruption in *Triobp^ΔEx9-10/ΔEx9-10^* mice (right). The data are representative of two *Triobp^ΔEx9-10/+^* and two *Triobp^ΔEx9-10/ΔEx9-10^* OPCs and two *Triobp^ΔEx9-10/+^* and three *Triobp^ΔEx9-10/ΔEx9-10^* DCs. **d** Faint fluorescent F-actin labeling could be revealed in OPCs in a 400nm-thick maximum intensity projection view at the level of the bottom edges of OHC cuticular plates. **e** A lateral view of microfilaments/microtubules shown in panel c. The insets show a smaller density of these structures in *Triobp^ΔEx9-10/ΔEx9-10^* OPCs. Scale bars: a, c, e – 1 μm; d – μmm.

Besides hair cell rootlets and cuticular plates, TRIOBP is also localized to the apical phalangeal projections of OPCs and DCs (Fig. 3b). The apical phalangeal microtubular projections are located deeper, at the level of the bottom edges of the hair cell cuticular plates. Transverse Z-projection of the 400nm-thick FIB-SEM volume at this level revealed distinct filament/microtubule structures in normal hearing *Triobp^ΔEx9-10/+^* OPCs (Fig. 3c, arrow) and dense actin patches in *Triobp^ΔEx9-10/+^* DCs (Fig. 3c, arrowhead). These structures were either disrupted or barely visible in homozygous deaf *Triobp^ΔEx9-10/ΔEx9-10^* littermates (Fig. 3c). It is worth mentioning that, besides microtubules [3], the apical projections of OPCs also contain actin filaments that can be revealed by fluorescently labelled phalloidin (Fig. 3d). Sagittal re-slicing of FIB-SEM volume parallel to the filament/microtubule structures in OPCs showed that they are present in both *Triobp^ΔEx9-10/+^* and *Triobp^ΔEx9-10/ΔEx9-10^* mice (Fig. 3e), but their density and regularity are impaired in mutant *Triobp^ΔEx9-10/ΔEx9-10^* OPCs (Fig. 3e, insets). Thus, TRIOBP-5 deficiency results in ultrastructural defects in OHCs, OPCs and DCs, which are the three major types of cells that form reticular lamina of the organ of Corti.

### Absence of TRIOBP-5 softens apical surfaces of supporting and hair cells and stereocilia rootlets

To investigate the role that TRIOBP isoforms exert on cochlear cell cytoskeletons and on the mechanical resistance of cochlear sensory epithelia, using PFT-AFM we measured the Young’s elastic modulus (E, stiffness) of the reticular lamina of wild-type and *Triobp* mutant mice. Both topography and mechanical mapping were obtained simultaneously by gently indenting the apical surface (<500nm) of living mouse organ of Corti explants at postnatal ages P5 to P6 (P5-P6) under physiologically relevant conditions. PFT-AFM nanomechanical mapping was conducted in the middle turn of the mouse organ of Corti along the radial axis of freshly dissected sensory epithelial explants.

The Young’s modulus of the localized nano-indented points was calculated. Nanoscale stiffness maps were generated using live P5-P6 organ of Corti of *Triobp^+/+^* (wild-type), heterozygous *Triobp^ΔEx9-10/+^,* and homozygous mutant *Triobp^ΔEx9-10/ΔEx9-10^* littermates (Fig. 4a and Supplementary Figure S3). On the apical surfaces of PCs and DCs, localized stiffnesses of the reticular lamina of mutant *Triobp^ΔEx9-10/ΔEx9-10^* mice were significantly reduced (p<0.0001) compared with that of normal hearing *Triobp^+/+^* littermates (Fig. 4b and Supplementary Table 1). Thus, an absence of TRIOBP-5 protein in the organ of Corti reticular lamina significantly diminished the apical stiffness of supporting cells. Unexpectedly, we observed that the local stiffness at the apical surfaces of normal hearing heterozygous *Triobp^ΔEx9-10/+^* mice is also significantly reduced to an intermediate level between wild-type *Triobp^+/+^* and *Triobp^ΔEx9-10/ΔEx9-10^* for PCs (p<0.01) and DCs (p<0.0001) arguing for a wild-type gene dose-dependent influence of TRIOBP-5 on the apical stiffness of supporting cells. Our quantitative ddPCR data described above indicate that heterozygous *Triobp^ΔEx9-10/+^* mice have reduced *Triobp5* mRNA expression that is approximately one half that of the wild-type level (Supplementary Fig. 1b). However, the reduction in stiffness of the reticular lamina in young heterozygote mice does not significantly harm later on the hearing ability as indicated by auditory brainstem response analyses at P14 [10]. Whether or not a heterozygous carrier of a recessive pathogenic variant of *TRIOBP*/*Triobp* is more susceptible to age-related hearing loss or the organ of Corti is at greater risk of damage from loud noise is now an important question to explore. Relevant to this issue, in a large human cohort, a GWAS study identified a missense variant of *TRIOBP* that affects only the *TRIOBP*-5 isoform that is associated with age-related hearing loss [18].

**Fig. 4.**
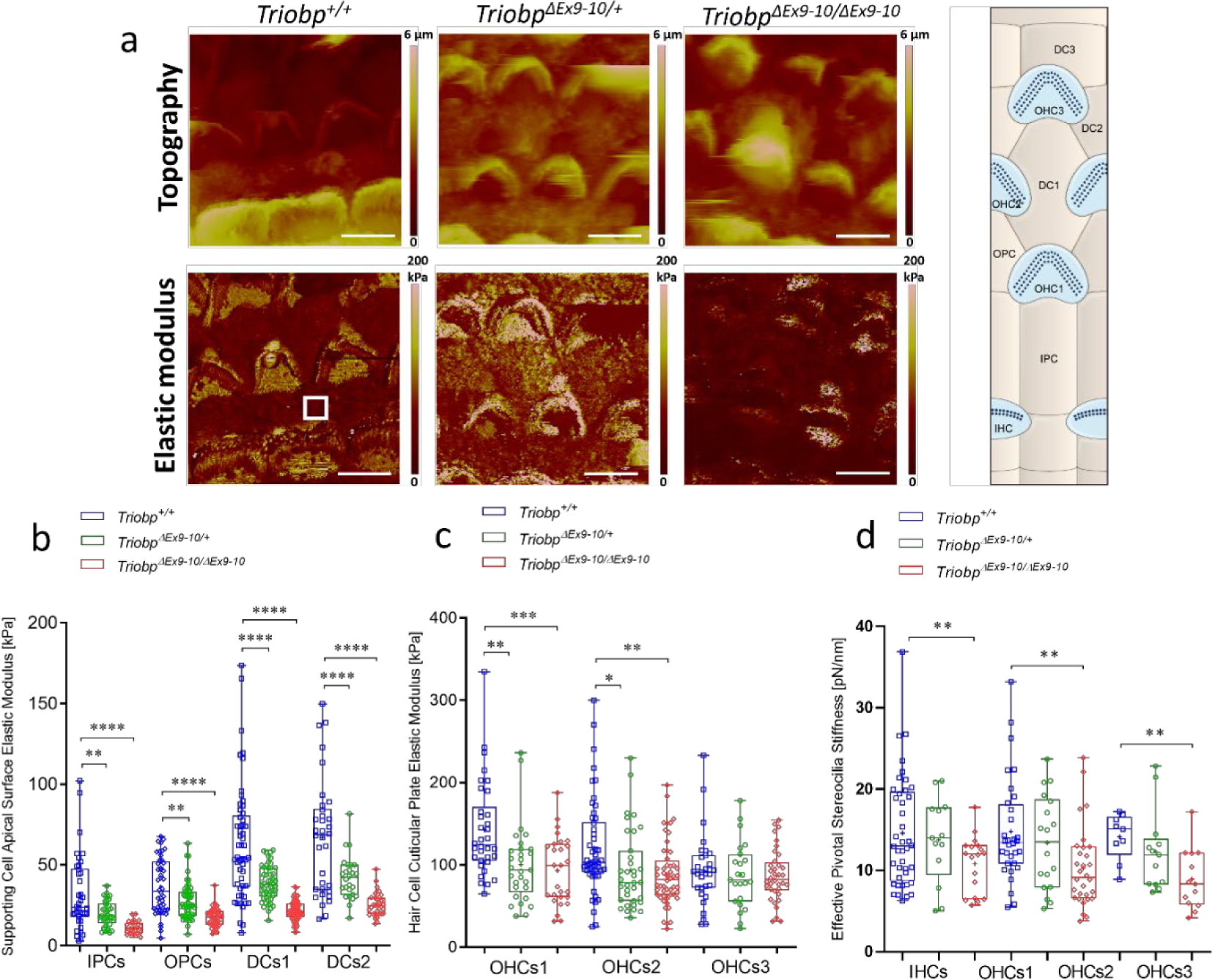
TRIOBP-5 deficiency results in a significant decrease in supporting cell apical surfaces stiffness and a decrease to a lesser degree in hair cell apical surface and stereocilia bundle stiffness. **a** Topography and stiffness maps of the organ of Corti explants for wild-type *Triobp^+/+^*, heterozygous *Triobp^ΔEx9-10/+^*, and homozygous *Triobp^ΔEx9-10/ΔEx9-10^* mice. The schematic illustrates the locations of the apical surfaces of supporting cells phalangeal process and the hair cells cuticular plate within the reticular lamina. **b** E values for apical processes of IPCs, OPCs, DCs row 1 (DCs1), and DCs row 2 (DCs2) of *Triobp^+/+^*, *Triobp^ΔEx9-10/+^*, and *Triobp^ΔEx9-10/ΔEx9-10^*, respectively. **c** E values for cuticular plates of three rows of OHCs including OHCs1, OHCs2, and OHCs3 of *Triobp^+/+^*, *Triobp^ΔEx9-10/+^*, and *Triobp^ΔEx9-10/ΔEx9-10^*, respectively. The E values of the reticular lamina were measured in regions of interest (white square box in *Triobp^+/+^* Young’s modulus image) overlaying the apical surface of supporting and hair cells. **d** Effective pivotal stereocilia stiffness values within hair bundles of three rows of OHCs. Data are represented as mean (kPa or pN/nm) ± standard deviation; Significant differences between conditions by unpaired two-tailed Student’s t-test with Welch’s correction indicated as, **** P˂0.0001, *** P˂0.001, **P˂0.01, and *P˂0.05; Scale bars are 5µm. See Supplementary table 1 for detailed statistical analyzes.

Notably, TRIOBP-5 is expressed in sensory epithelium supporting cells and localized to cytoskeletal filamentous structures within the DCs apical processes and apical surfaces of IPCs and OPCs [10]. In *Triobp^ΔEx9-10/ΔEx9-10^* mice, a TRIOBP-5-specific antibody signal is absent from supporting cells [10]. Therefore, a reduction in the apical stiffness of supporting cells likely results from an alteration of these developing cytoskeletal filamentous structures in supporting cells of mutant *Triobp^ΔEx9-10/ΔEx9-10^* mice, which could later affect sound-induced intracochlear stimulations and hearing function [27, 29, 30].

In addition, the local stiffnesses of the cuticular plates in outer hair cells of mutant *Triobp^ΔEx9-10/ΔEx9-10^* mice are significantly reduced (varying from p<0.001-0.1) as compared with wild-type *Triobp^+/+^* (Fig. 4c and Supplementary Table 1). Interestingly, similar to supporting cell mechanical behavior, we observed that the local stiffness of OHCs cuticular plates in normal hearing heterozygous *Triobp^ΔEx9-10/+^* mice is also reduced compared to wild-type *Triobp^+/+^* (varying from p<0.01-0.1). ddPCR assays confirmed an approximately 60-percentage reduction in TRIOBP-5 mRNA expression in heterozygous *Triobp^ΔEx9-10/+^* OHCs compared to wild-type during postnatal development at P6 (Supplementary Fig. 1b). Thus, TRIOBP-5 deficiency in the reticular lamina of the organ of Corti is associated with a decrease in apical stiffness of supporting cells and hair cell cuticular plates.

Next, we investigated the role of TRIOBP-5 in the pivotal stiffness of stereocilia within outer hair cell bundles using PFT-AFM performing stereocilia nano-deflections in the excitatory (positive) direction at physiologically relevant deflection magnitudes of <200nm [31]. For OHC bundle stiffness, the local effective stiffness of mutant *Triobp^ΔEx9-10/ΔEx9-10^* mice deficient for *TRIOBP-5* is significantly reduced as compared with the wild-type *Triobp^+/+^* (p<0.01) (Fig. 4d and Supplementary Table 1). Thus, TRIOBP-5 deficiency in the organ of Corti is sufficient to cause a change in the effective pivotal stiffness of stereocilia bundles. These observations agree with our previous results using fluid-jet deflection of the stereocilia bundle [10]. Deflections of the mutant *Triobp^ΔEx9-10/ΔEx9-10^* mouse hair bundles under the fluid-jet pressure were increased more than that of *Triobp^+/+^* hair bundles at the same low stimulus intensities of <200nm [10]. Thus, with similar low stimulus intensities, the fluid-jet and PFT-AFM techniques both showed an increase in pivotal flexibility of stereocilia bundles of TRIOBP-5 isoform-specific knockout mice as compared to the wild-type *Triobp^+/+^* mouse.

### Simultaneous absence of TRIOBP-4 and TRIOBP-5 further reduce stiffness of the reticular lamina and hair bundles

Using PFT-AFM, we generated topography and stiffness maps of P5-P6 live organ of Corti explants (middle turn) from wild-type *Triobp^+/+^*, heterozygotes *Triobp^ΔEx8/+^,* and homozygous mutant *Triobp^ΔEx8/ΔEx8^* littermates (Fig. 5a and Supplementary Figure S4). On apical surfaces of PCs and DCs of *Triobp^ΔEx8/ΔEx8^* mice, deficient for both TRIOBP-4 and TRIOBP-5, local stiffness of the reticular lamina was significantly reduced (P<0.001) as compared with *Triobp^+/+^* mice (Fig. 5b and Supplementary Table 2). The local stiffness of the PC and DC apical surfaces of heterozygous *Triobp^ΔEx8/+^* normal hearing mice is also reduced compared to wild-type PCs (p<0.05) and DCs (P<0.0001). In comparison with changes reported for a TRIOBP-5 deficiency alone, the combined absence of TRIOBP-4 and TRIOBP-5 shows a greater decrease of the local axial stiffness of supporting cells within the reticular lamina. Since TRIOBP-4 and TRIOBP-5 were previously reported to localize to the cytoskeletal filamentous structures within the apical surfaces of the supporting PCs and DCs [10, 26], and taking into account our stiffness measurement data, we conclude that both TRIOBP-4 and TRIOBP-5 isoforms contribute to the mechanical resistance of supporting cells within the reticular lamina. Moreover, the local stiffness of the reticular lamina at the cuticular plates of hair cells of *Triobp^ΔEx8/ΔEx8^* mice is significantly reduced (varying from P<0.0001-0.05) compared with wild-type mice (Fig. 5c and Supplementary Table 2). The simultaneous deficiency for TRIOBP-4 and TRIOBP-5 in the organ of Corti resulted in a more profound decrease in the mechanical resilience of the cuticular plate of OHCs. This suggests that TRIOBP-4 and TRIOBP-5 are both reinforcing hair cell cuticular plates and the absence of both TRIOBP isoforms significantly soften the cuticular plate.

**Fig. 5.**
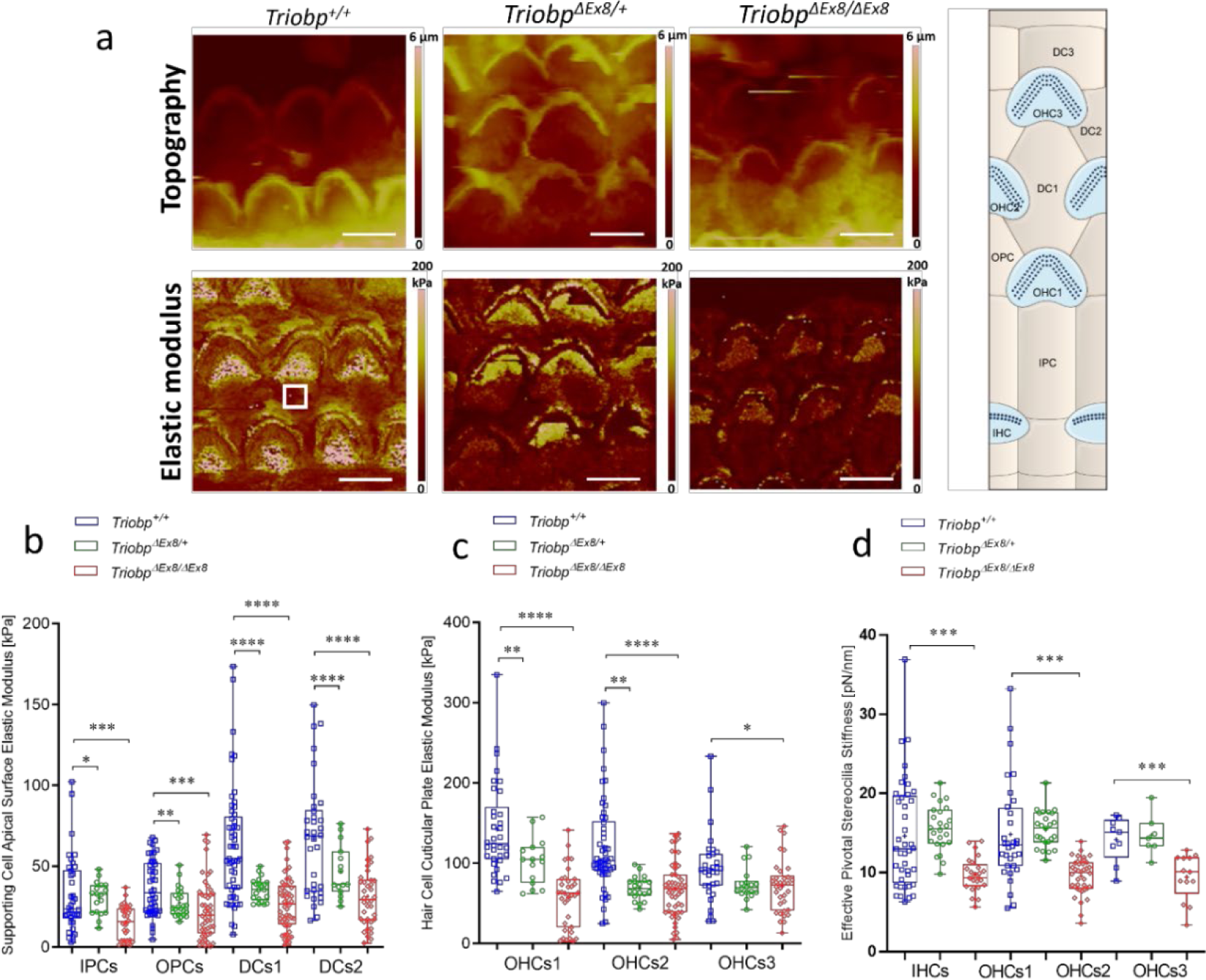
Deficiency of both TRIOBP-4 and TRIOBP-5 proteins results in significant decreases in stiffness of supporting and hair cell apical surfaces and stereocilia bundles. **a** Topography and stiffness maps of the organ of Corti explants for wild-type *Triobp^+/+^*, heterozygous *Triobp^ΔEx8/+^*, and homozygous *Triobp^ΔEx8/ΔEx8^* mice. The schematic illustrates the locations of the apical surfaces of supporting cells phalangeal process and the hair cells cuticular plate within the reticular lamina. **b** E values for apical processes of IPCs, OPCs, DCs row 1 (DCs1), and DCs row 2 (DCs2) of *Triobp^+/+^*, *Triobp^ΔEx8/+^*, and *Triobp^ΔEx8/ΔEx8^*, respectively. **c** E values for cuticular plates of three rows of OHCs including OHCs1, OHCs2 and OHCs3 of *Triobp^+/+^*, *Triobp^ΔEx8/+^* and *Triobp^ΔEx8/ΔEx8^*, respectively. The E values of the reticular lamina were measured in regions of interest (white square box in *Triobp^+/+^* Young’s modulus image) overlaying the apical surface of supporting and hair cells. **d** Effective pivotal stereocilia stiffness values within hair bundles of three rows of OHCs. Note, *Triobp^+/+^* wild-type control results are the same for Figures 4 and 5 since both mice strains used in this study are on the same C57BL/6 background. Data are represented as mean (kPa or pN/nm) ± standard deviation; Significant differences between conditions by unpaired two-tailed Student’s t-test with Welch’s correction indicated as, **** P˂0.0001, *** P˂0.001, **P˂0.01, and *P˂0.05; Scale bars are 5µm. See Supplementary table 2 for detailed statistical analyzes.

To investigate the effects of a deficiency of both TRIOBP-4 and TRIOBP-5 on the mechanical resilience of stereocilia rootlets, we analyzed the mechanical changes of local points within the hair bundle. Local effective stiffness of the OHCs hair bundles revealed that *Triobp^ΔEx8/ΔEx8^* stereocilia are much more flexible as compared to *Triobp^+/+^* and heterozygous *Triobp^ΔEx8/+^* stereocilia (P<0.001) (Fig. 5d and Supplementary Table 2). Compared to the observations of a TRIOBP-5 only deficiency, the simultaneous loss of TRIOBP-4 and TRIOBP-5 isoforms (TRIOBP-4/5) resulted in a more significant increase in the mechanical compliance of the stereocilia bundles. These results agree with an increased fragility of stereocilia from *Triobp^ΔEx8/ΔEx8^* mice that we reported previously using fluid-jet stimulations [9]. At the same low stimuli intensities, the extent of the nano-deflection of the hair bundles in the TRIOBP-4/5 isoform-specific knockout mouse was significantly larger as compared with that measured in the wild-type. Furthermore, our previous immunofluorescence observations revealed a restricted localization of TRIOBP-5 to the rootlet segments embedded within a hair cell cuticular plate. By comparison, TRIOBP-4 is localized along the entire length of rootlets and also diffusely distributed within the OHC cuticular plates [10]. Thus, a simultaneous loss of both TRIOBP-4 and TRIOBP-5 significantly impacts resilience of the F-actin cytoskeletal structures of the hair cell cuticular plates and stereocilia bundles to a greater degree as compared to a loss of only TRIOBP-5, which in mouse results in progressive hearing loss with reduced severity compared to the profound congenital deafness of the *Triobp* mutant mouse deficient for both TRIOBP-4 and TRIOBP-5 proteins [9, 10].

### Hair cells and supporting cells have radial stiffness gradients of opposing orientations

High-resolution PFT-AFM nanomechanical mapping revealed unexpected radial stiffness gradients in the mouse organ of Corti. PFT-AFM analyses allowed us to visualize the spatial changes in stiffness along the radial axis of live organ of Corti explants. The local stiffness of the reticular lamina at the apical surfaces of nonsensory supporting cells increases from PCs toward DCs. However, the local stiffness of the reticular lamina at the cuticular plates of sensory hair cells declines from the first toward the third row of OHCs. Thus, there are complex and bidirectional radial stiffness gradients in the organ of Corti reticular lamina consisting of oppositely oriented radial gradients for hair cells versus supporting cells (Fig. 6a). Perhaps, supracellular epithelial tensional homeostasis [32] of the cochlear sensory tissue is achieved by balancing the mechanical properties between supporting cells and hair cells.

**Fig. 6.**
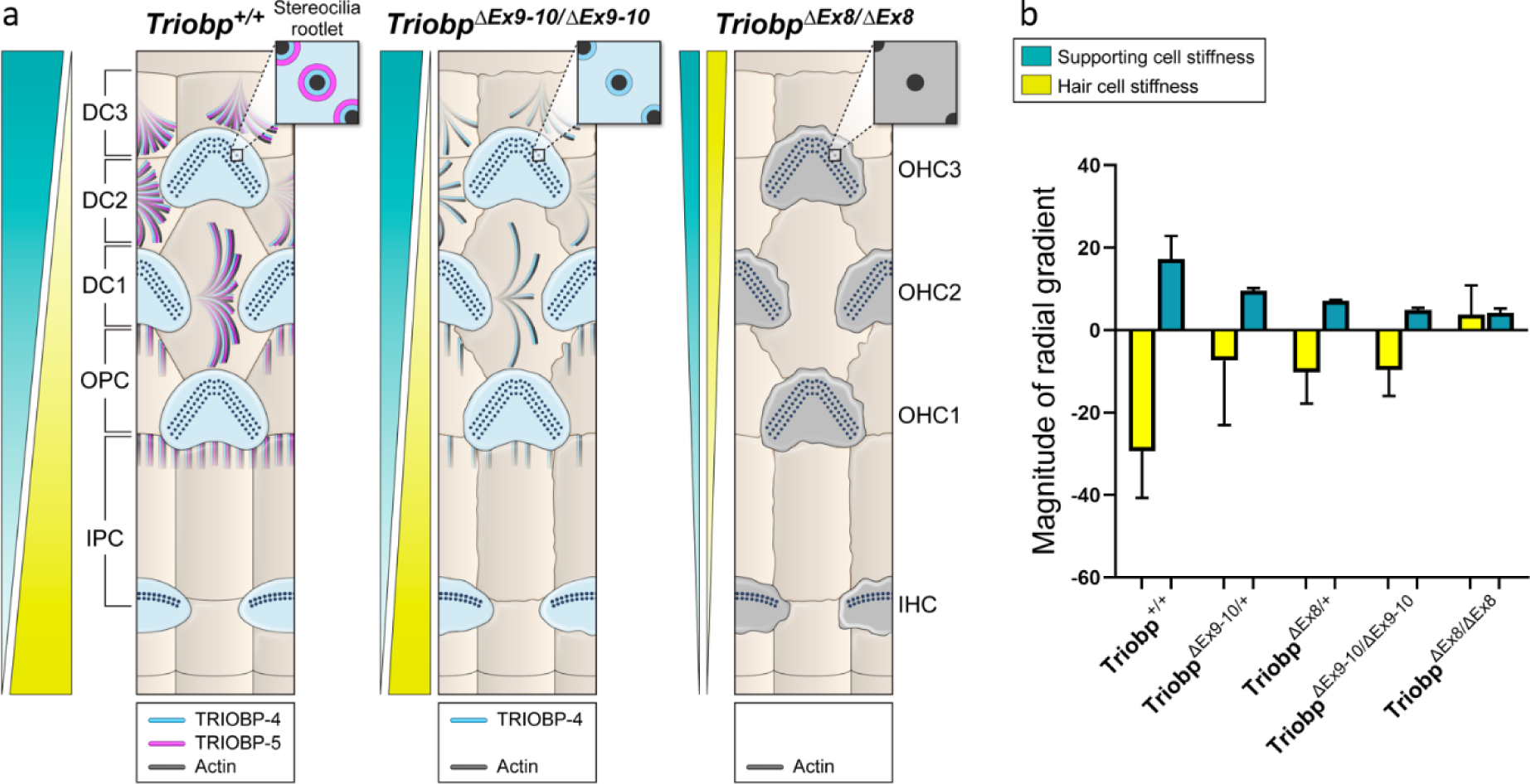
Summary model describing opposing radial stiffness gradients in the reticular lamina of the organ of Corti. **a** Illustration of TRIOBP-4 and TRIOBP-5 distribution in the stereocilia bundles, cuticular plate of hair cells and apical processes of supporting cells of wild- type (*Triobp^+/+^*), *Triobp^ΔEx9-10/ΔEx9-10^* homozygous deficient for TRIOBP-5, and *Triobp^ΔEx8/ΔEx8^* homozygous deficient for both TRIOBP-4 and TRIOBP-5 (TRIOBP-4/5) and also demonstrating radial stiffness gradient in the organ of Corti reticular lamina of each one. **b** Average magnitudes of radial gradients in Young’s modulus values against cell position within the reticular lamina (slope; E/radial position) of supporting and hair cells for *Triobp^+/+^*, heterozygous *Triobp^ΔEx9-10/+^,* and homozygous *Triobp^ΔEx9-10/ΔEx9-10^* mice as well as heterozygous *Triobp^ΔEx8/+^,* and homozygous *Triobp^ΔEx8/ΔEx8^* mice. Note that when TRIOBP-4 and TRIOBP-5 isoforms are both absent the reticular lamina radial stiffness gradients become unidirectional. Data are represented as mean ± standard error of the mean. See Supplementary table 3x for detailed statistical analyzes.

### Absence of both TRIOBP-4 and TRIOBP-5 diminishes reticular lamina bidirectional radial stiffness gradients

High-resolution PFT-AFM nanomechanical mapping was also utilized to monitor the radial stiffness gradient in the mouse organ of Corti reticular lamina of the wild-type *Triobp^+/+^*, heterozygous, and homozygous mutant littermate mice. We first investigated the reticular lamina radial stiffness gradients behavior when only TRIOBP-5 was absent. For all genotypes (*Triobp^+/+^*, *Triobp^ΔEx9-10/+^,* and *Triobp^ΔEx9-10/ΔEx9-10^*), the local stiffness of the reticular lamina at the apical surfaces of supporting cells increases from the PCs toward the DCs (positive gradient). However, the magnitude of the radial stiffness gradient of supporting cell apical surfaces was reduced in the *Triobp* mutants compared with the wild-type *Triobp^+/+^* mouse (Fig. 6b and Supplementary Table 3). Furthermore, for all conditions, the local stiffness at the cuticular plates of sensory hair cells decreases from the first toward the third row of OHCs showing a negative gradient. Similarly, the magnitude of the radial stiffness gradient of the hair cell cuticular plates was significantly reduced (P<0.01) as compared to the *Triobp^+/+^* mouse (Fig. 6b and Supplementary Table 3). Therefore, the absence of TRIOBP-5 reduces the reticular lamina radial stiffness gradients, although these stiffness gradients are not completely abolished.

Similar to *Triobp-5* only deficient mice, in the case of Triobp-4 and Triobp-5 deficiencies (*Triobp^+/+^, Triobp^ΔEx8/+^,* and *Triobp^ΔEx8/ΔEx8^* mice), the local stiffness of the reticular lamina at the apical surfaces of supporting cells increases from the PCs toward the DCs (positive gradient). In addition, the radial stiffness gradient magnitude of supporting cell apical surfaces was significantly reduced in the *Triobp^ΔEx8/+^* and *Triobp^ΔEx8/ΔEx8^* mutants compared with wild-type *Triobp^+/+^* (Fig. 6b and Supplementary Table 3). Interestingly, in heterozygous *Triobp^ΔEx8/+^* mice, the magnitude of the radial stiffness gradient of the outer hair cell cuticular plates is reduced but maintains its orientation similar to that of wild-type *Triobp^+/+^*, while in homozygous *Triobp^Δex8/Δex8^* mice, the gradient orientation was reversed (Fig. 6b and Supplementary Table 3). Altogether, absence of TRIOBP-4 and TRIOBP-5 or TRIOBP-5 alone significantly reduces the magnitudes of the reticular lamina radial stiffness gradient of supporting and outer hair cells, whereas the absence of both TRIOBP isoforms significantly alters the normal behavior (magnitude and orientation) of the radial stiffness gradient of hair cells. Therefore, both TRIOBP-4 and TRIOBP-5 are individually critical for the maintenance of optimal tissue-level reticular lamina radial stiffness gradients and their absence compromises the cochlear mechanical properties, which in turn may contribute to the pathophysiology of TRIOBP-deficient mouse and phenotype of human DFNB28 deafness.

## Discussion

In the organ of Corti, mechanical interactions among neighboring OHCs depend on intricate relationships of the adjacent mosaic of supporting cells (PCs, OPCs, and DCs), which collaborate to form the reticular lamina [33-35]. No previous study has mapped the cochlear reticular lamina at nanoscale resolution due to the intrinsic low spatial resolution of previous biophysical approaches using glass micro-pipettes [23, 24], fluid jets [25], or standard atomic force microscopy methods [26]. In this study, using a nanoscale resolution PFT-AFM method, we measured the mechanical properties of the spatially heterogeneous apical surface of the cochlear sensorial reticular lamina, including the individual hair cells and supporting cells apical surfaces and revealed previously unknown radial gradients of stiffness with opposite orientations when supporting cell local stiffnesses are compared with hair cells.

Using two *Triobp* deaf mouse models that recapitulate human recessively inherited deafness DFNB28 [9, 10], we demonstrated essential roles in mechanical homeostasis of the F-actin-associated TRIOBP-4 and TRIOBP-5 proteins in inner ear tissue. Our findings indicate that the absence of both *Triobp* isoforms significantly reduce the stiffness of supporting cell apical surfaces, hair cell cuticular plates and stereociliary bundles. Furthermore, our data also show the existence of cochlear reticular lamina composite radial stiffness gradients, which decreases from the first toward the third row of OHCs in the cuticular plates of the hair cells and increases from the second row of DCs to the IPCs in the apical surfaces of the supporting cells. These radial gradient magnitudes of stiffness are significantly diminished in *Triobp* mutant mice with greater reductions observed when TRIOBP-4 and TRIOBP-5 proteins are both absent as compared to loss of TRIOBP-5 only.

Microtubules and bundled actin filaments together are essential for the mechanical properties of the apical surfaces of sensory hair cells and supporting cells [36]. This interplay was studied using some molecular mediators such as latrunculin A, jasplakinolide, blebbistatin, taxol, and nocodazole [27]. Here, our observations argue for the importance of the F-actin- bunding proteins TRIOBP-4 and TRIOBP-5 for the characteristic mechanical properties of the cytoskeletal components of hair cells and supporting cells and for their overall apical surface stiffness. Recently, GAS2 (growth-arrest-specific 2), a protein with microtubule and F-actin binding domains was shown to be expressed in pillar and Deiters’ cells co-localizing specifically with microtubules. Absence of GAS2 significantly reduced the stiffness of the supporting cell phalangeal processes [36]. These observations raise additional questions as to the individual functions of other cytoskeleton-associated proteins of hair cells and supporting cells such as tropomyosin, spectrin, β-actin, γ-actin, espin, prestin, and tubulin and the effects of pathogenic variants in the corresponding genes on the cellular structures that might also contribute to the radial stiffness gradients we report here.

A longitudinal gradient along the length of the cochlear reticular lamina have been reported for frequency-detection of sound, for hair cell and supporting cell morphology [3], and for the stiffness of organ of Corti individual sensory cells [37]. For example, the relative stiffness of pillar cells is greater in the basal region of the cochlea where high frequency sound is detected, while stiffness decreases toward the apex of the cochlea where low frequency sound is detected. These surface mechanical properties of sensory and nonsensory cells have been attributed to changes in the cytoskeletal architecture [27]. In addition, a longitudinal gradient in the length of stereocilia and the OHC bodies increases from base-to-apex [37]. Interestingly, previous studies have also shown longitudinal and radial gradients of compositional, structural, and mechanical properties in the basilar membrane and in the acellular tectorial membrane that covers organ of Corti hair cells [38, 39]. The overall structure and collagen fibril orientation and thickness of the tectorial membrane varies along the cochlea, which are hypothesized to underlie a decreasing basal-to-apical gradient in tectorial membrane stiffness in the region overlying hair cell stereocilia [38]. The evidence for and implications of a longitudinal gradient along the cochlea [40] and a radial gradient [41] in tectorial membrane elasticity were reported and associated with a radial gradient in collagen fibril density [41, 42]. In addition, different mechanical properties of the basilar membrane, such as its increasing stiffness towards the base, are graded along the length of the cochlea [39], although the factors that lead to the gradient in its mechanical properties are still unclear. Since longitudinal and radial gradients in mechanical properties have been previously shown in the tectorial and basilar membranes, we hypothesized that the organ of Corti reticular lamina may also possess a distinct and intricate radial gradient. We further hypothesize that existence of a radial stiffness gradient in the reticular lamina could help maintain the organ of Corti intricate sensitivity and frequency discrimination by providing a passive cochlear mechanical filtering that defines which outer hair cell would enhance the auditory vibration stimuli of particular frequencies. To the best of our knowledge, our study is the first to demonstrate a complex radial mechanical stiffness gradient in the organ of Corti reticular lamina.

The organ of Corti mechanical properties regulate cochlear amplification [43, 44]. Specifically, two mechanical conditions are responsible for location-dependent amplification of sound of high to low frequencies from base-to-apex of the cochlea. First, the phase of the outer hair cell somatic electromotility force varies along the cochlear length. Second, the local stiffness of the organ of Corti varies along the cochlear length. However, the mechanisms for similar gradients in the radial direction of the organ of Corti remain unclear, although a radial morphology and gene expression asymmetries in the organ of Corti have been shown previously in the sensory hair cells of both adult and developing tissues [33]. For instance, there are prominent longitudinal and radial gradients of the expression of prestin, a motor protein in the lateral wall of outer hair cells responsible for electromotility [45]. Prestin immunostaining intensity increases in outer cells from base-to-apex and from first to the third row of OHCs [46].

A radial gradient in the mechanical properties of the organ of Corti is supported by computational simulations [47] and *in vivo* imaging [48]. Sasmal and Grosh developed a mathematical model showing that the reticular lamina may indeed has radial passive and active mechanical properties [47]. Radial tuning in the reticular lamina of the mouse organ of Corti has also been reported by *in vivo* volumetric optical coherence tomography vibrometry measurements [48]. Interestingly, this radial tuning of the reticular lamina is detected in a dead cochlea, when OHC no longer produce prestin-driven amplification of the electromotility force, and in the detached tectorial membrane mutants of TECTA, suggesting that the radial tuning of the reticular lamina is derived mainly from passive mechanical properties, including apical surface passive tension, stiffness, and viscosity, consistent with our discovered radial gradients in the measured passive mechanical property (Young’s modulus). Additionally, radial tuning (passive mechanical properties) of the reticular lamina is not imparted, controlled or modified by the tectorial membrane, justifying our preparations for measuring the apical stiffness of organ of Corti explants. Khanna and Hao conducted experiments on the apical turn of living guinea pig cochleae and reported a significant increase in detected vibration amplitude and phase shift in the radial direction of hair cells from IHCs toward the third row of OHCs [49], suggesting a softening of hair cells in the radial direction, consistent with our measurements. They also reported a significantly larger magnitude of detected amplitude of Hensen’s cells compared to that of the hair cells, suggesting that supporting cells are significantly softer when compared to hair cells, an observation that is also consistent with our measurements.

The bidirectional radial stiffness gradients discovered in this study provides new experimental evidence of reticular lamina mechanical coordination to maximize hair bundle deflection, thus improving hearing sensitivity, amplification, and tone suppression. Any radial change in stiffness is significant because it would allow the outer hair cells from the 1^st^ toward the 3^rd^ row to absorb high frequency energy. This behavior is similar to the base to apex cochlear partition ability to absorb high frequency energy changes in fluid pressure, which declines towards the apex [50]. Interestingly, a bidirectional change in radial stiffness with hair cells and supporting cells having different stiffness behaviors suggest diverse mechanisms of sound frequency discrimination possibly allowing a balanced force transmission and simple radial shear within the reticular laminar. In other words, these opposite gradients would cancel each other to make the overall reticular lamina stiffness more uniform. Otherwise, the longer OHCs of the third row, which are less stiff than shorter OHCs of the first row, would have a much lower rotational stiffness and would be less sensitive. Future studies in mouse with conditionally ablated TRIOBP isoform expression only in supporting cells are needed to test the hypothesis that the radial stiffness gradient and changes in local supporting cell stiffness that we observed in *Triobp* mutant mice are required for normal hearing.

In conclusion, our atomic force microscopy-based nanomechanical maps revealed complex, bidirectional radial stiffness gradients in the reticular lamina of the mouse cochlea. The magnitude of these radial gradients requires the expression of TRIOBP in hair cells and supporting cells. We also discovered hitherto unreported spatial mechanical gradients within the reticular lamina, which are different for inner ear sensory and non-sensory cells. Previously we showed that the presence of both TRIOBP-4 and TRIOBP-5 is necessary for stereocilia rootlet formation and architecture, and that hair cells with stereocilia lacking rootlets will subsequently die, causing deafness [9, 10]. Here, we demonstrated that the loss of these two TRIOBP isoforms also compromises optimal tissue-level radial stiffness gradients of opposite orientation and mechanical homeostasis of the auditory sensory epithelium in the organ of Corti.

## Methods

### Mouse models and genotyping

Wild-type *Triobp^+/+^*, heterozygous *Triobp^ΔEx9-10/+^,* and homozygous *Triobp^ΔEx9-10/ΔEx9-10^* mice as well as heterozygous *Triobp^ΔEx8/+^* and homozygous *Triobp^ΔEx8/ΔEx8^* mice were generated and genotyped as described previously [9, 10].

### In situ hybridization using RNAscope probes

*In situ* hybridizations were performed using RNAscope Multiplex Fluorescent V2 assay (Advanced Cell Diagnostics (ACD), Newark, CA) with following probes. The Probe-Mm-Triobp-O1 that targeted the region from nucleotide 387 to 1288 (NM_001039155.1) was used to detect both *Triobp-4* and *Triobp-5* mRNAs. The Probe-Mm-Triobp-O2-C3 targeted the region from nucleotide 3860 to 4773 of NM_138579.4 and was used to detect only *Triobp*-5 mRNA. The Probe-Mm-Triobp-O3 targeted the region from nucleotide 154 to 1170 of NM_001024716.1 and was used to detect both *Triobp*-1 and *Triobp-5*. As a positive control, Probe-Mm-Myo7a-C2 targeted the region from nucleotide 1365 to 2453, NM_001256081.1 of *Myo7a* mRNA was used to label IHCs and OHCs. Cochleae from C57BL/6J wild-type mice at P6 and P14 were fixed overnight at 4°C in 4% PFA (Electron Microscopy Sciences, Hatfield, PA) in 1x PBS. Fixed P14 cochleae were decalcified in 10% EDTA for 5 days. Then, cochleae were cryopreserved in 15% sucrose in 1x PBS overnight at 4°C and then in 30% sucrose in 1X PBS overnight at 4°C. Each cochlea was embedded and frozen in Super Cryoembedding Medium (Section-Lab, Hiroshima, Japan). Frozen cochleae were sectioned at a thickness of 12µm using a CM3050S cryostat microtome (Leica, Vienna, Austria). Images were taken with an LSM 880 confocal microscope equipped with 63x and 40x objectives (Carl Zeiss Microscopy, Thornwood, NY).

### Droplet digital PCR

In order to evaluate the expression profile of different *Triobp* mRNA isoforms, whole inner ears were dissected from wild-type, heterozygous and homozygous *Triobp^ΔEx9-10^* mice at P6. Total RNA was extracted using Trizol (Invitrogen, USA) followed by first-strand cDNA synthesis using a SuperScript™ RT-PCR kit (Invitrogen, USA) according to manufacturer instructions. TaqMan probes for different isoforms of *Triobp* were designed (Supplementary Fig. 1a), *Triobp-1* primers spanned the junction between exon 1b, which is unique for *Triobp1* mRNA, to exon 13, with probe designed over the exon junction [Triobp_1- FWD GGGAAGGGCTGGAGCTA; Triobp_1-REV AGATTGACATCCATCCTTTCTTGA; Triobp_1-PRB /56FAM/CATGACGCC/ZEN/CGATCTGCTCAACT/3IABkFQ/]. Whereas for detecting *Triobp-5*, the primers were designed spanning the junction from exon 8 to exon 9 and the TaqMan probe designed over the junction [Triobp_5-FWD AGGTGGAGCGCCTCTTC; Triobp_5-REV AAGTTGGCTCTGAGCTTGG; Triobp_5-PRB /56-FAM/AAGAGCGCA/ZEN/GGAAATCGGAGGC/3IABkFQ/]. To detect cDNA synthesized from*Triobp-4* mRNA, a forward primer was designed in exon 8, and the reverse primer was located in sequence of the exon 8 UTR unique only to *Triobp-4* [Triobp_4-FWD GCAAGAGCGCAGGTGAG; Triobp_4-REV CAGCAGGGCTTGAACTCTT; Triobp_4-PRB/56-FAM/TGGAACTTT/ZEN/CCAACTGTTCTCCTCCC/3IABkFQ/] (Supplementary Fig. 1a).

The expression of the *Triobp* alternative splice isoforms was quantified using a QX200 ddPCR System (Bio-Rad, USA) according to the manufacturer’s instructions. Briefly, a 20μL ddPCR reaction comprised of 2x ddPCR Supermix for Probes (no dUTP) (Bio-Rad, USA), 20x TaqMan assay primer-probe mix and cDNA sample. Droplets were generated using the Automated Droplet Generator (Bio-Rad, USA). The PCR plate was subsequently heat-sealed with pierceable foil using a PX1 PCR plate sealer (Bio-Rad, USA) and then amplified in a Veriti thermal cycler (Applied Biosystems, USA) using the following amplification cycling conditions, with initial enzyme activation at 95°C for 10min, followed by 40 cycles of denaturation at 94°C for 30 sec and an annealing/extension step at 60°C for 1 min. Final enzyme inactivation at 98°C for 10min and held at 4°C until the next step. After amplification, the 96-well plate was fixed in a plate holder and placed into the QX200 Droplet Reader (Bio-Rad, USA) and droplets from each well of the plate were read automatically. QuantaSoft analysis software (Bio-Rad, USA) was used to analyze ddPCR data and for the quantification of the target molecule per 1μl of PCR reaction.

### Electron tomography with Focused Ion Beam serial sectioning and Scanning Electron Microscopy imaging (FIB-SEM)

Cochleae from heterozygous (*Triobp^ΔEx9-10/+^*) and homozygous (*Triobp^ΔEx9-10/ΔEx9-10^*) mice at postnatal day 6 (P6) were extracted from temporal bones and gently perfused through the oval window with a solution containing 2.5% glutaraldehyde, 2% paraformaldehyde in 0.1M cacodylate buffer, pH=7.4 (Electron Microscopy Science, cat# 15960-01), and supplemented with 2mM CaCl2 and 1% Tannic acid (Electron Microscopy Sciences, cat# 21710). The cochleae were kept in this fixative overnight at 4°C, and then the fixative was diluted ∼1:5 with 0.1M cacodylate buffer and samples were shipped to the University of Kentucky. Then, the organs of Corti were dissected in distilled water with the tectorial membrane kept intact and high-pressure frozen with Leica EM ICE high-pressure freezer. The frozen samples were transferred to Leica EM AFS2 freeze substitution machine and kept in methanol with 1% uranyl acetate at -90°C for 33 hours and then slowly warmed at about 4°C per hour to -45°C. Samples were then washed with fresh 100% methanol for 24 hours at -45°C to fully replace uranyl acetate. Then the methanol was gradually replaced with Lowicryl HM-20 resin (monostep embedding kit, Electron Microscopy Sciences, cat# 14345): 50% Lowicryl (2 hours), 75% Lowicryl (overnight ∼21 hours), and 100% Lowicryl (overnight ∼24 hours). To ensure that there is no residual methanol, the samples were additionally incubated with fresh 100% Lowicryl for 1 hour. Then, samples were transferred into flat mold with 100% Lowicryl, incubated for additional 24 hours, and polymerized with UV light at -45°C for 27 hours, at 0°C for additional 40 hours, and at 20°C for 48 hours. The resin blocks were trimmed (Leica EM TRIM2) to reach a desired sample distance of 20-50µm from the upper surface of the block. Samples were then sputter coated with 25nm of platinum (Electron Microscopy Science, EMS150T ES) and serial sectioned with a focused ion beam at a 20nm step size and imaged in “Slice and View” mode with a backscattered electron detector using the FEI Helios 660 Nanolab system.

### Analysis of FIB-SEM stacks

Raw FIB-SEM stacks typically contained several hundreds of images with 6144×4096 pixels with a vortex size of ∼2×2×20nm. They were registered either with a custom MATLAB script or with SIFT registration plugin in ImageJ-2 (Fiji, National Institutes of Health). After 3D median filtering, the stacks were rotated with Interactive Stack Rotation plugin in Fiji in such a way that the field of view became parallel to the feature of interest (upper surface of the cuticular plate for hair cells or reticular lamina for supporting cells). These rotated stacks were used to construct top median Z-projection views of the cell of interest. To reveal microtubules/microfilaments within the outer pillar cells, the stacks were further oriented in such a way that microfilaments/microtubules would align with X-axis and then the stack was rotated 90 degrees along the X-axes, providing a lateral view of these structures. Thus, a starting orientation of the field of view parallel to the cuticular plate of hair cells (or reticular lamina in supporting cells) ensures the match of orientations of resulting Z-projections between the samples.

### Immunostaining

P14 wild-type *Triobp^+/+^* and homozygous *Triobp^ΔEx9-10/ΔEx9-10^* mice were used for immunostaining of the organ of Corti with TRIOBP-5 antibody as previously described [9, 10]. Briefly, organs of Corti were fixed in 2% paraformaldehyde for 30 min at room temperature (RT), microdissected and permeabilized in 0.2% Triton X-100 in 1X PBS for 15min at RT, washed in 1X PBS and incubate in blocking solution (2% BSA and 5% normal goat serum in 1X PBS) overnight at 4^0^C. Next day the samples were transferred to the custom-made primary rabbit polyclonal anti-TRIOBP-5 antibody [9, 10] diluted in blocking solution 1:200 to obtain 5µg/ml concentration and incubated for 2 hours at RT, followed by 3 × 10min washes in 1X PBS. Then samples were incubated in secondary anti-rabbit Alexa Fluor 488 conjugated antibody (Thermo Fisher, Cat # R37118) at 1:400 dilution together with Rhodamine-phalloidin (Thermo Fisher, Cat # R415) at 1:100 dilution in blocking solution for 30min at RT, followed by 3 × 10min washes in 1X PBS and mounted on glass slides using ProLong Gold Antifade mountant (Thermo Fisher, Cat # P36934). Prepared slides were kept in a slide box at 4^0^C overnight and then examined using LSM 780 Zeiss confocal microscope equipped with 63x, 1.4 N.A. oil immersion objective.

### Organ of Corti explant cultures

Wild-type *Triobp^+/+^*, heterozygous *Triobp^ΔEx9-10/+^*, and homozygous *Triobp^ΔEx9-10/ΔEx9-10^* mice, as well as heterozygous *Triobp^ΔEx8/+^* and homozygous *Triobp^ΔEx8/ΔEx8^* mice were used for explant culture experiments. Three P6 pups of each genotype were sacrificed by decapitation according to the National Institutes of Health Guidelines for Care and Use of Laboratory Animals. The bullae were removed from mouse temporal bones and submerged in cold Leibovitz media (L-15, 21083-027, Life Technologies). The organ of Corti sensory epithelium was microdissected from the cochleae, the tectorial membrane was removed using 27-gauge needle and the entire spiral of the organ of Corti was plated on a glass-bottom petri dish (HBST-5040, Willco Wells) precoated with 10μl of Cell-Tak (Corning) to immobilize it, and immersed in Leibovitz media (L-15, 21083-027, Life Technologies). The organ of Corti culture was incubated at 37⁰C and 5% CO2 for 15-30min to achieve firm attachment of the specimen before the PFT-AFM measurements.

### PFT-AFM imaging

PeakForce Tapping mode (PFT-AFM) images were obtained using a Bruker BioScope Catalyst AFM system (Bruker) mounted on an inverted Zeiss Axiovert 200M optical microscope equipped with a 40x objective (0.95 NA, Plan-Apochromat, Zeiss) and a confocal laser scanning microscope (LSM 510 META, Zeiss). During PFT-AFM experiments, the organ of Corti sensory epithelium explants were maintained at 37°C using a Bruker heated stage and scanned using a probe appropriate for live cell imaging with a tip height of 17μm, controlled tip radius of 65nm, and opening angle of 15° (PFQNM-LC, Bruker). The cantilever probes used had spring constant values ranging between 0.06–0.08N/m and were pre-calibrated by the Bruker AFM system using the thermal tune method. Probes were replaced for each new experiment or more frequently as needed. The images were collected at a scan resolution of 128 × 128 pixels and a scan rate of 0.5Hz. Each tissue scan lasted ∼10 minutes. The driving frequency was set at 500Hz and the drive amplitude was set up to 850nm. The peak force setpoint was kept between 800pN and 1.2nN. The Young’s modulus stiffness maps of the samples were analyzed and extracted via the NanoScope Analysis software (Bruker) using the Sneddon’s contact mechanics model in which F_Sneddon_ = (8Etanα⁄3 π)δ^2^, where *F* is the applied force, *α* is the tip half-opening angle, and *δ* is the sample mean indentation [51].

### Point stiffness of stereocilia

The PFT-AFM experiments produced continuous series of force-distance curves throughout the tested cochlear samples. Discrete raw force-distance curves across the hair cell bundles were acquired and the noncontact virtual deflection tilt (generated by hydrodynamic drag) of the baseline was corrected using NanoScope Analysis software (Bruker). Then, individual force-distance curves were quantitatively analyzed using a custom-made MATLAB-based data processing code to determine the apparent point (localized) stiffness, available upon request. The point stiffness was determined by performing a linear curve fitting process. Thus, the point stiffness is the slope of the force curve. Note, that the PFT- AFM-based hair bundle deflection from the probe-hair bundle contact point was limited to ∼200nm for the force-distance curves to calculate the pointed apparent stiffness. Our measurements was limited to 200nm of stereocilia deflection because in normal hearing the physiological deflections of hair bundles are between 80nm to 200nm [31].

### Statistical Analysis

All experiments were repeated at least 3 times. Data in Figs 4 and 5 are shown in Box-and-whisker plots. Bars show mean ± standard deviation (SD). Data in Fig. 6 are shown as mean ± standard error of the mean (SEM). Student’s t tests of independent variables (2-tailed t-test with Welch’s correction) were used to determine statistical significance and the asterisks indicate the level of statistical significance (*P<0.05, **P<0.01, ***P < 0.001, and ****P<0.0001).

### Study approval

The experiments on mice were conducted according to the National Institutes of Health Guidelines for Care and Use of Laboratory Animals. All experimental procedures were approved by the NINDS/NIDCD Animal Care and Use Committee (ACUC) at the National Institutes of Health (protocol 1263-20 to T.B.F).

## Authors Contributions

I.A.B., T.B.F., and A.X.C.R. conceived the project. H.B., I.A.B., R.Y., R.T., S.H., G.F., and A.X.C.R. conducted the experiments. H.B., I.A.B., T.B.F., and A.X.C.R. co-wrote the manuscript and all authors discussed and edited the manuscript.

## Acknowledgements

The authors acknowledge Mr. Alan Hoofring (NIH’s Medical Arts Design Section) for help with Illustrations (Figures 1 and 6). In addition, the authors thank Ms. Sherly Michel (NIDCD) and Mr. Pat Diers (NIDCD) for help with animal care and genotyping and Dr. Kuni Iwasa (NIDCD) and Dr. Benjamin Perrin (Indiana University - Purdue University Indianapolis) for carefully reading the manuscript and providing critical inputs. A.X.C.R. and this work was supported by the National Institutes of Health (NIH) Distinguished Scholars Program and the NIH Intramural Research Program of the National Institute of Biomedical Imaging and Bioengineering (grant # ZIA EB000094). This research was supported (in part) by the Intramural Research Program of the NIH, NIDCD DC000039 to T.B.F. S.K. was supported by JSPS KAKENHI grant 20K09687 and G.F. by NIDCD/NIH (R01DC014658 and S10OD025130). The electron microscopy was performed at the University of Kentucky Electron Microscopy Center, which belongs to the National Science Foundation NNCI Kentucky Multiscale Manufacturing and Nano Integration Node, supported by ECCS-1542174.

---

**Supplementary Fig. 1.**
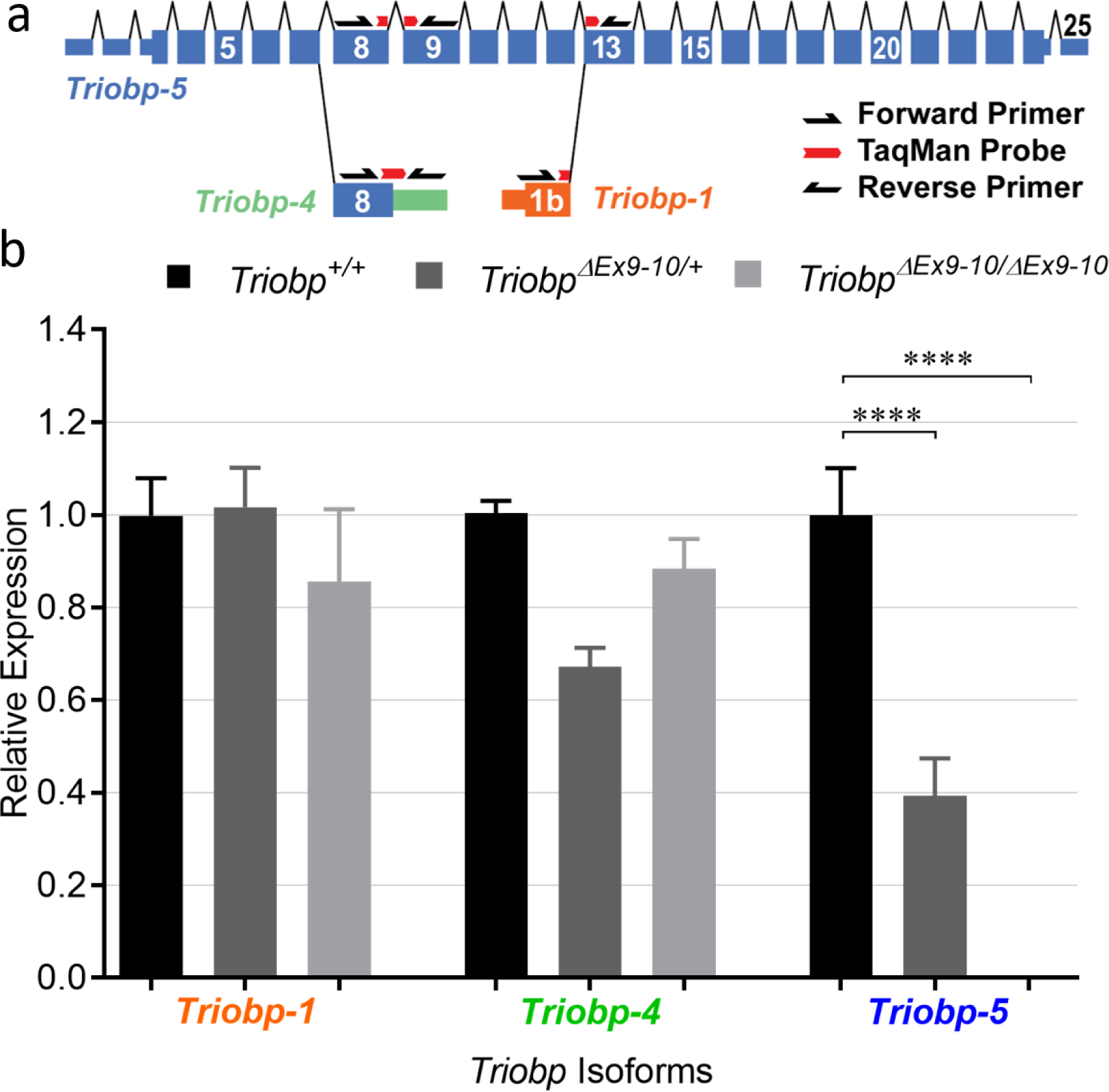
Expression profile of different isoforms in the *Triobp^ΔEx9-10^* mouse inner ear at P6. **a** TaqMan probes were designed to quantify different isoforms of Triobp mRNA. For Triobp-1 primers were designed that flanked the exon junction between exon 1b (orange), which is unique to Triobp1 mRNA, and exon 13, with probe designed over the exon junction. Whereas, for Triobp-5, the primers were designed flanking the exon 8 - 9 exon junction, with the probe designed over the spanning junction. To detect cDNA synthesized from Triobp-4 mRNA, a forward primer was designed in the coding region of exon 8. The reverse primer was located in sequence of the 3’ UTR of exon 8 unique only to Triobp-4 (Green). **b** Expression levels of three *Triobp* mRNA isoforms (*Triobp-1*, *Triobp-4* and *Triobp-5*) which were measured using ddPCR. There did not appear to be statistically significant difference in expression levels of either *Triobp-1 or Triobp-4* isoforms among the three genotypes. However, the *Triobp-5* expression level was reduced by approximately 40% in heterozygous *Triobp^ΔEx9-10/+^* mice, a genotype that ablates 2 exons of *Triobp-5* in one genomic copy of the Triobp gene. No mRNA expression of *Triobp-5* was detected in homozygous *Triobp^ΔEx9-10/ ΔEx9-10^* mice. Data are represented as mean ± standard error of mean; significant differences between conditions by unpaired two-tailed Student’s t-test with Welch’s correction indicated as **** P<0.0001.

**Supplementary Fig. 2 (related to Figure 2).**
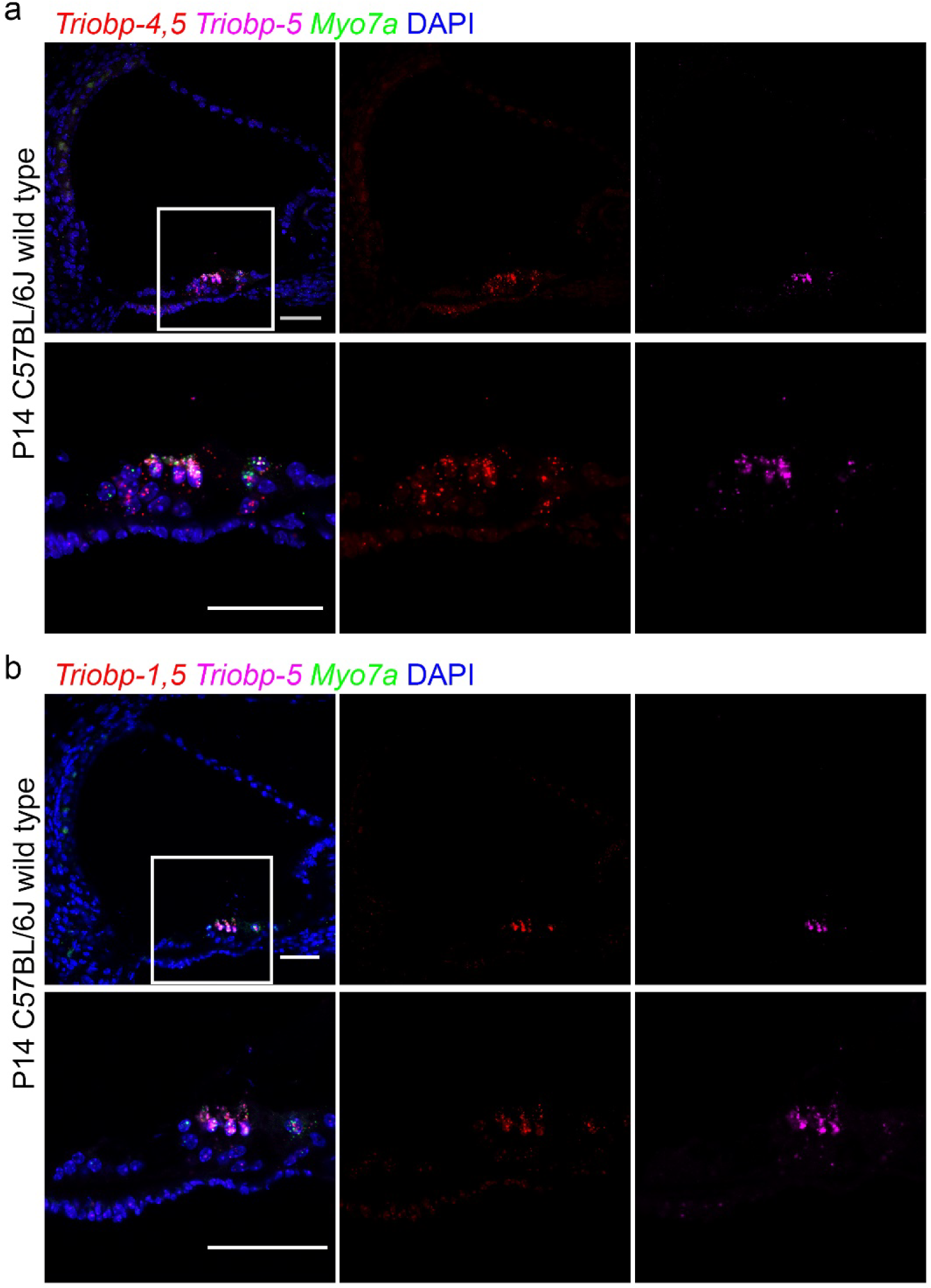
RNAscope probe *in situ* hybridization in P14 wild-type mouse cochlea. **a** Expression of *Triobp-4 and Triobp-5* mRNAs (*Triobp-4/5*) (red, Probe-Mm-Triobp-O1), *Triobp-5* only mRNA (magenta, Probe-Mm-Triobp-O2-C3) and *Myo7a* mRNA (green, Probe-Mm-Myo7a-C2). *Triobp-4/5* mRNA (red) is expressed in inner and outer hair cells and supporting cells. Whereas mRNA of *Triobp-5* alone is expressed mainly in OHCs. **b** Expression of both *Triobp-1 and Triobp-5* mRNA (*Triobp-1/5*) (red, Probe-Mm-Triobp-O3), *Triobp-5* only mRNA (magenta, Probe-Mm-Triobp-O2-C3) and *Myo7a* mRNA (green, Probe-Mm-Myo7a-C2). *Triobp-1/5* mRNA was detected mainly in hair cells. Scale bars are 50µm.

**Supplementary Fig. 3 (related to Figure 4).**
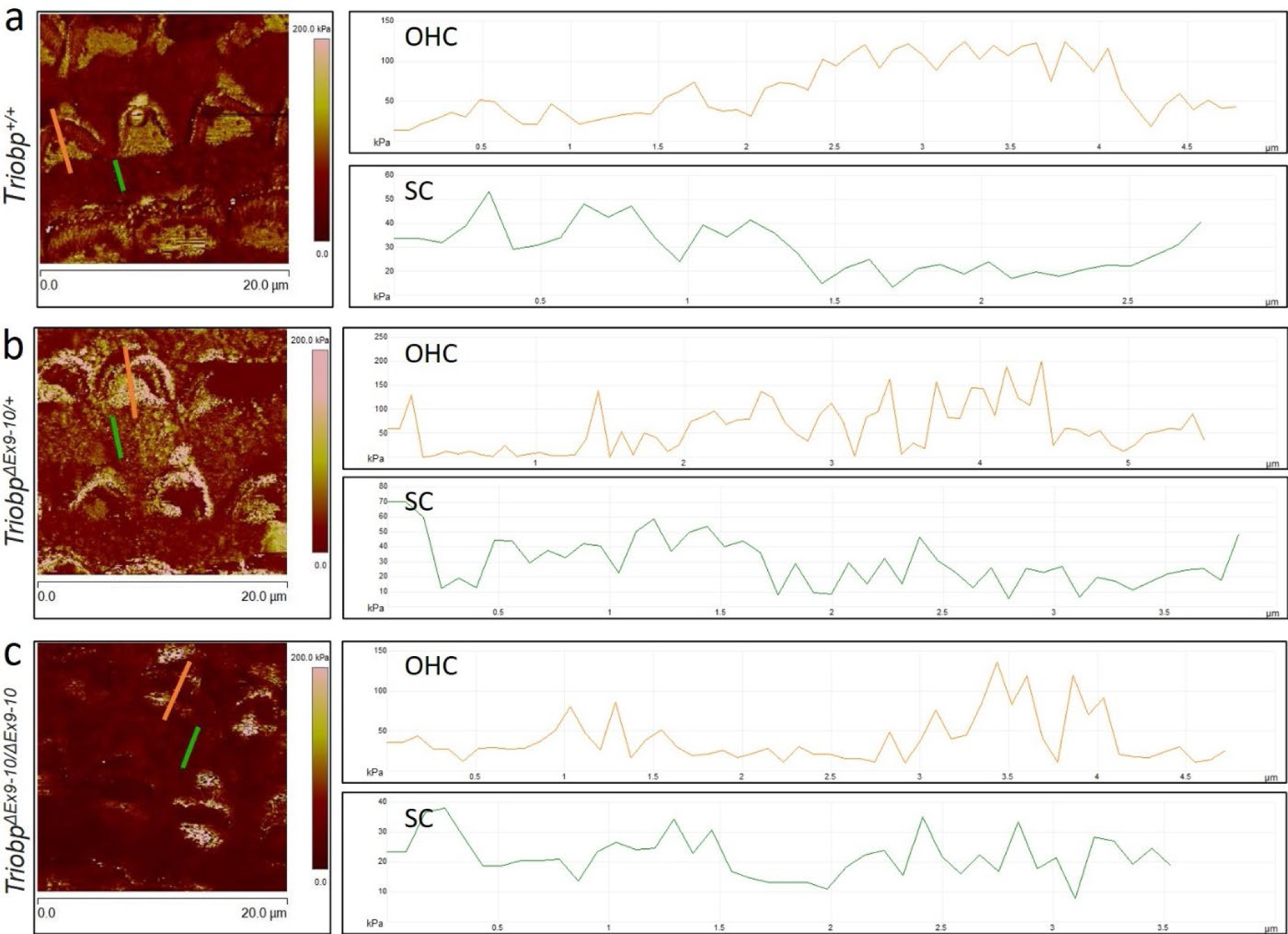
Extracted line profiles from TRIOBP-5 deficient mice cochlear explants PFT-AFM maps shows nanometer scale variations of apical surface Young’s modulus on outer hair cells and supporting cells. Stiffness maps and line profiles of the organ of Corti explants for wild-type *Triobp^+/+^* **(a)**, heterozygous *Triobp^ΔEx9-10/+^* **(b)**, and homozygous *Triobp^ΔEx9-10/ΔEx9-10^* mice **(b)**. The orange and green line profiles represent stiffness data of an outer hair cell and a supporting cell, respectively.

**Supplementary Fig. 4 (related to Figure 5).**
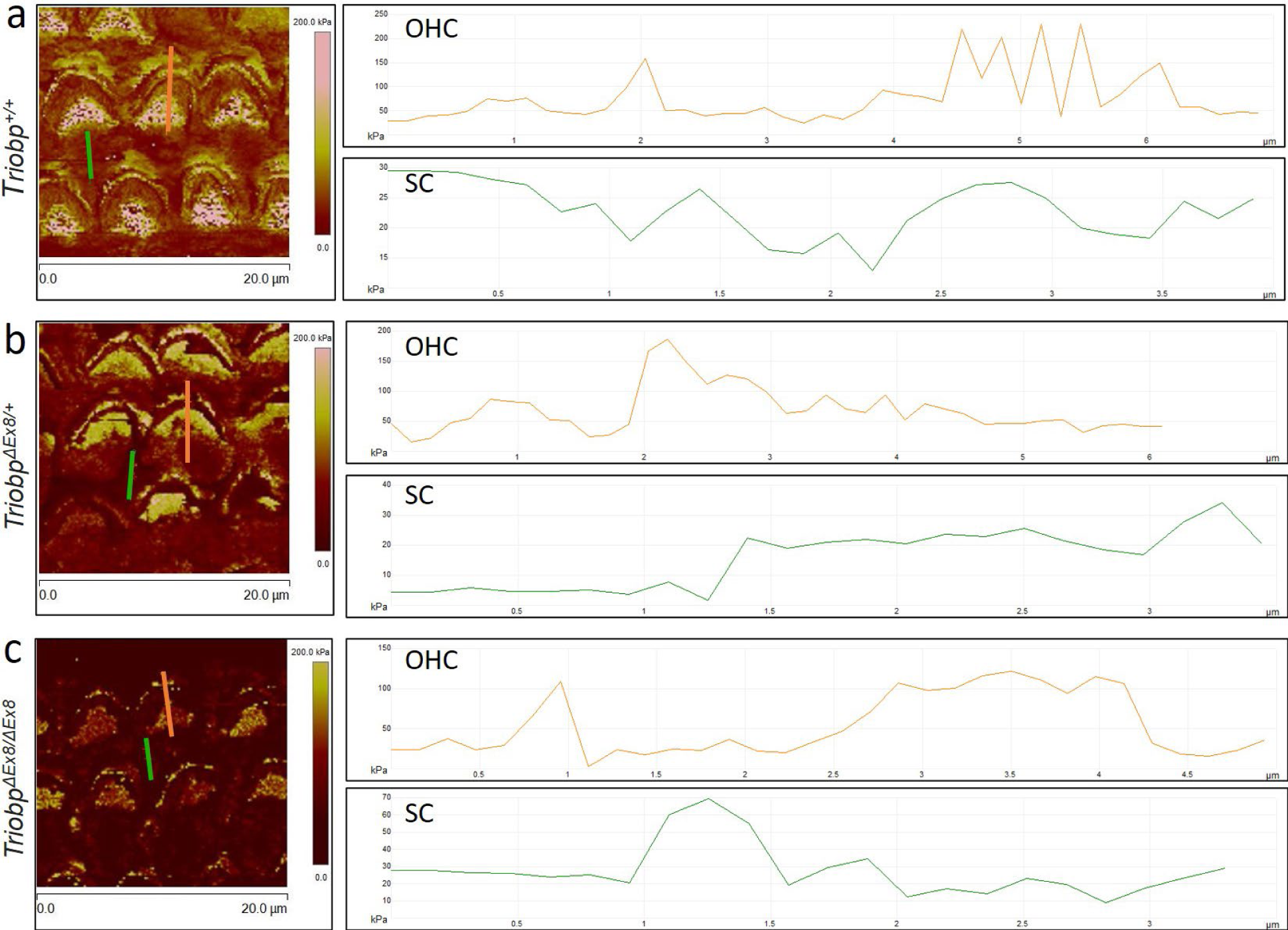
Extracted line profiles from TRIOBP-4/5 deficient mice cochlear explants PFT-AFM maps shows nanometer scale variations of apical surface Young’s modulus on outer hair cells and supporting cells. Stiffness maps and line profiles of the organ of Corti explants for wild-type *Triobp^+/+^* **(a)**, heterozygous *Triobp^ΔEx8/+^* **(b)**, and homozygous *Triobp^ΔEx8/ΔEx8^* mice **(b)**. The orange and green line profiles represent stiffness data of an outer hair cell and a supporting cell, respectively.

**Supplementary Table 1 (related to Figure 4).**
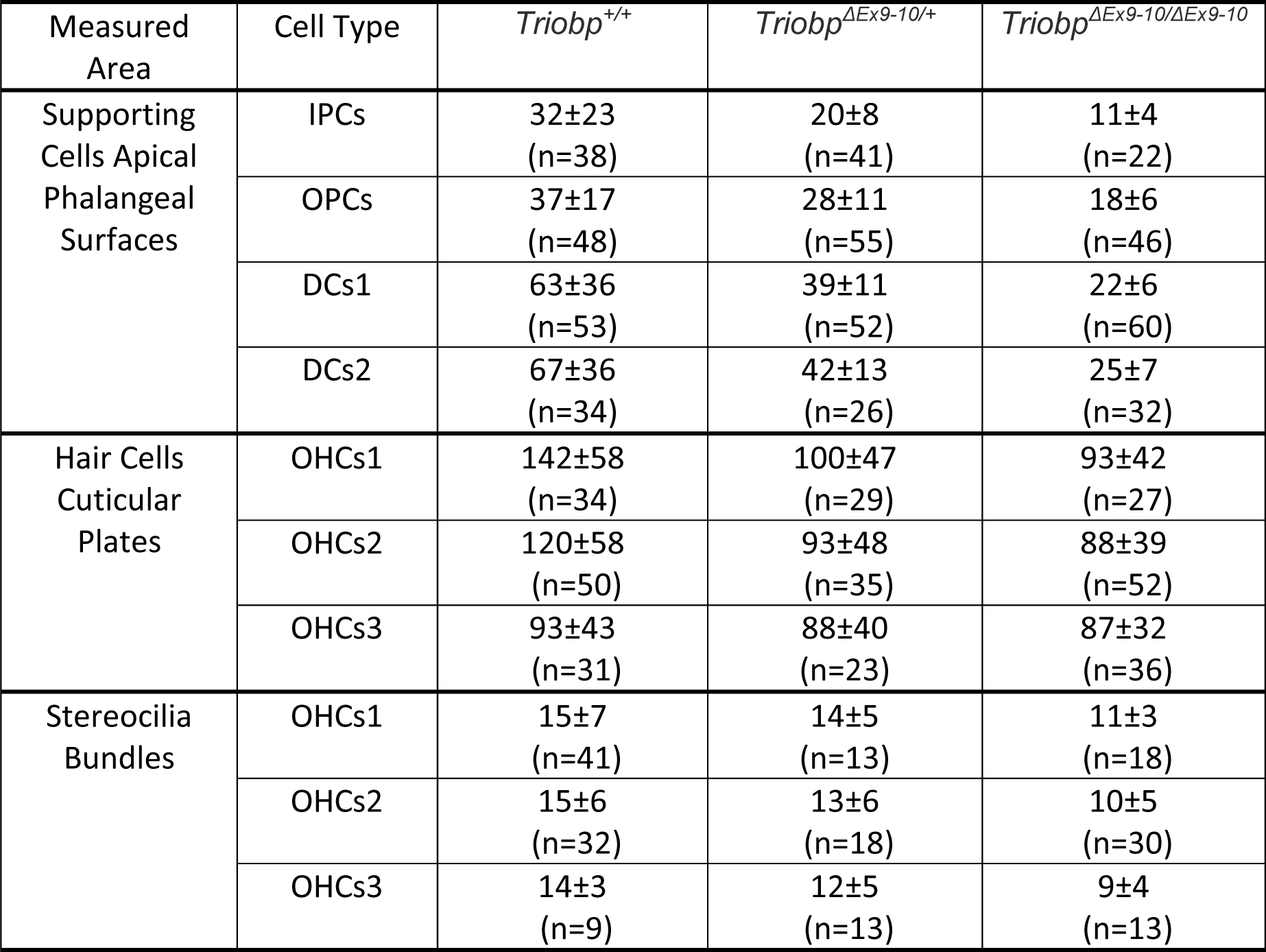
Summarized measurements for the Young’s modulus (kPa) of the organ of Corti middle turn apical surface of supporting cells and cuticular plate of hair cells and stiffness (slope, pN/nm) of stereocilia within hair bundles obtained from live P5-P6 explants for wild-type *Triobp^+/+^*, heterozygous *Triobp^ΔEx9-10/+^*, and homozygous *_Triobp_ΔEx9-10/ΔEx9-10* mice. *mean (kPa) ± standard deviation and n is the number of cells. IPCs, Inner pillar cells; OPCs, Outer pillar cells; DC, Deiters’ cells; OHCs, Outer hair cells. Represented data were acquired in replicates for *Triobp^+/+^* wild-type mice (4 animals), *Triobp^ΔEx9-10/+^* heterozygous mice (5 animals), and *Triobp^ΔEx9-10/ΔEx9-10^* homozygous mice (5 animals).

| Measured Area | Cell Type | *Triobp*<sup>+/+</sup> | *Triobp*<sup>ΔEx9-10/+</sup> | *Triobp*<sup>ΔEx9-10/ΔEx9-10</sup> |
| --- | --- | --- | --- | --- |
| Supporting Cells Apical Phalangeal Surfaces | IPCs | 32±23<br>(n=38) | 20±8<br>(n=41) | 11±4<br>(n=22) |
|  | OPCs | 37±17<br>(n=48) | 28±11<br>(n=55) | 18±6<br>(n=46) |
|  | DCs1 | 63±36<br>(n=53) | 39±11<br>(n=52) | 22±6<br>(n=60) |
|  | DCs2 | 67±36<br>(n=34) | 42±13<br>(n=26) | 25±7<br>(n=32) |
| Hair Cells Cuticular Plates | OHCs1 | 142±58<br>(n=34) | 100±47<br>(n=29) | 93±42<br>(n=27) |
|  | OHCs2 | 120±58<br>(n=50) | 93±48<br>(n=35) | 88±39<br>(n=52) |
|  | OHCs3 | 93±43<br>(n=31) | 88±40<br>(n=23) | 87±32<br>(n=36) |
| Stereocilia Bundles | OHCs1 | 15±7<br>(n=41) | 14±5<br>(n=13) | 11±3<br>(n=18) |
|  | OHCs2 | 15±6<br>(n=32) | 13±6<br>(n=18) | 10±5<br>(n=30) |
|  | OHCs3 | 14±3<br>(n=9) | 12±5<br>(n=13) | 9±4<br>(n=13) |

\*mean (kPa) ± standard deviation and n is the number of cells. IPCs, Inner pillar cells; OPCs, Outer pillar cells; DC, Deiters' cells; OHCs, Outer hair cells. Represented data were acquired in replicates for *Triobp*<sup>+/+</sup> wild-type mice (4 animals), *Triobp* <sup>$\Delta$ Ex9-10/+</sup> heterozygous mice (5 animals), and *Triobp* <sup>$\Delta$ Ex9-10/ ΔEx9-10</sup> homozygous mice (5 animals).

**Supplementary Table 2 (related to Figure 5).**
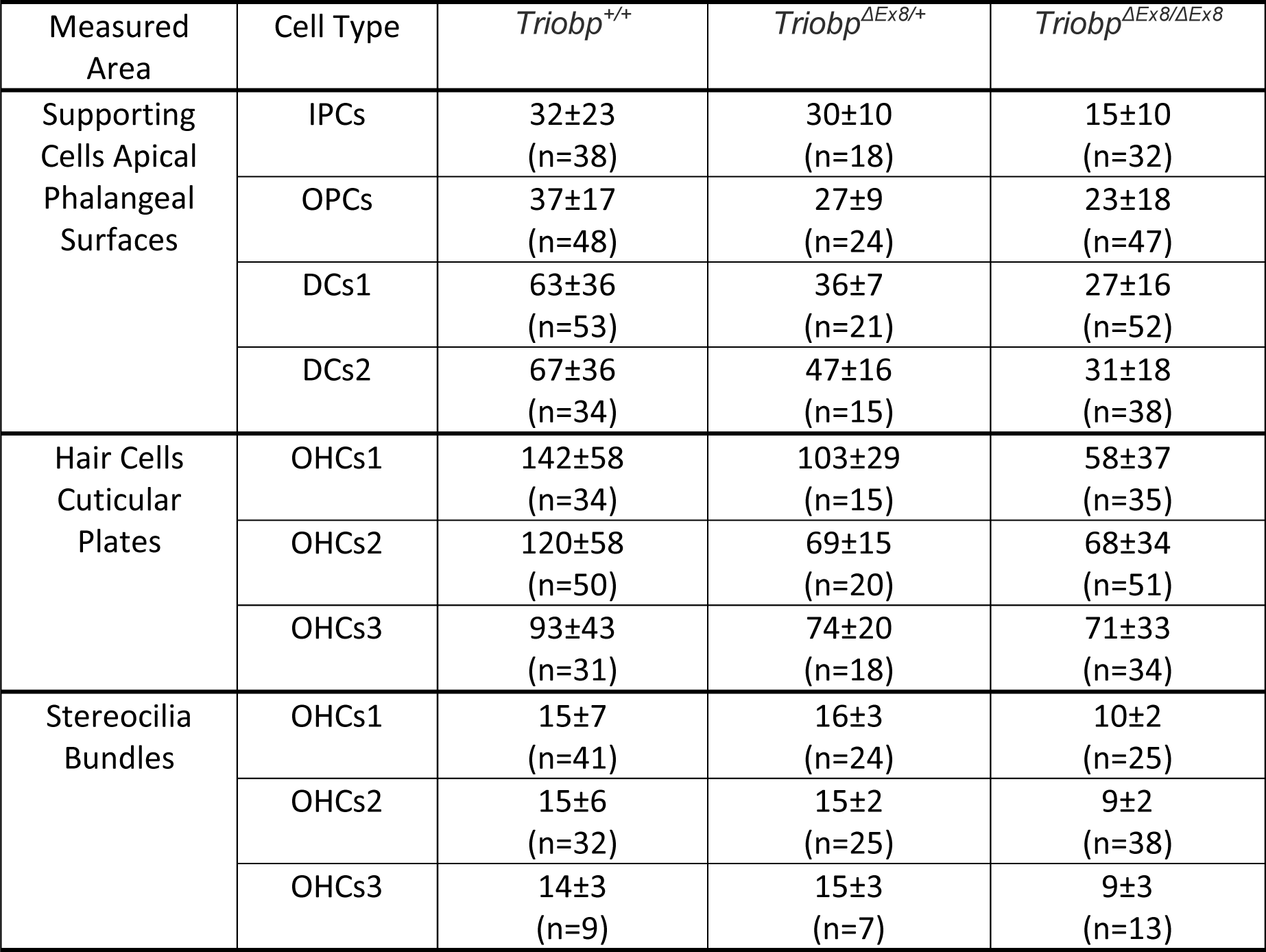
Summarized measurements for the Young’s modulus (kPa) of the organ of Corti middle turn apical surface of supporting cells and cuticular plate of hair cells and stiffness (slope, pN/nm) of stereocilia within hair bundles obtained from live P5-P6 explants for wild-type *Triobp^+/+^*, heterozygous *Triobp^ΔEx8/+^*, and homozygous *Triobp^ΔEx8/ΔEx8^* mice. *mean (kPa) ± standard deviation and n is the number of cells. IPCs, Inner pillar cells; OPCs, Outer pillar cells; DC, Deiters’ cells; OHCs, Outer hair cells. Represented data were acquired in replicates for *Triobp^+/+^* wild-type mice (4 animals), *Triobp^ΔEx8/+^* heterozygous mice 5 animals), and *Triobp^ΔEx8/ΔEx8^* homozygous mice (5 animals).

| Measured Area | Cell Type | *Triobp*<sup>+/+</sup> | *Triobp*<sup>ΔEx8/+</sup> | *Triobp*<sup>ΔEx8/ΔEx8</sup> |
| --- | --- | --- | --- | --- |
| Supporting Cells Apical Phalangeal Surfaces | IPCs | 32±23<br>(n=38) | 30±10<br>(n=18) | 15±10<br>(n=32) |
|  | OPCs | 37±17<br>(n=48) | 27±9<br>(n=24) | 23±18<br>(n=47) |
|  | DCs1 | 63±36<br>(n=53) | 36±7<br>(n=21) | 27±16<br>(n=52) |
|  | DCs2 | 67±36<br>(n=34) | 47±16<br>(n=15) | 31±18<br>(n=38) |
| Hair Cells Cuticular Plates | OHCs1 | 142±58<br>(n=34) | 103±29<br>(n=15) | 58±37<br>(n=35) |
|  | OHCs2 | 120±58<br>(n=50) | 69±15<br>(n=20) | 68±34<br>(n=51) |
|  | OHCs3 | 93±43<br>(n=31) | 74±20<br>(n=18) | 71±33<br>(n=34) |
| Stereocilia Bundles | OHCs1 | 15±7<br>(n=41) | 16±3<br>(n=24) | 10±2<br>(n=25) |
|  | OHCs2 | 15±6<br>(n=32) | 15±2<br>(n=25) | 9±2<br>(n=38) |
|  | OHCs3 | 14±3<br>(n=9) | 15±3<br>(n=7) | 9±3<br>(n=13) |

\*mean (kPa) ± standard deviation and n is the number of cells. IPCs, Inner pillar cells; OPCs, Outer pillar cells; DC, Deiters' cells; OHCs, Outer hair cells. Represented data were acquired in replicates for *Triobp*<sup>+/+</sup> wild-type mice (4 animals), *Triobp* <sup>$\Delta$ Ex8/+</sup> heterozygous mice (5 animals), and *Triobp* <sup>$\Delta$ Ex8/ ΔEx8</sup> homozygous mice (5 animals).

**Supplementary Table 3.** **(related to Figure 6)** Summary of radial gradients in Young’s modulus values against cell positions within the organ of Corti reticular lamina (slope; E/radial position) of supporting and hair cells obtained from live P5-P6 explants for wild-type *Triobp^+/+^*, heterozygous *Triobp^ΔEx9-10/+^,* and *Triobp^ΔEx8/+^*, homozygous *Triobp^ΔEx9-10/ΔEx9-10^*, and *Triobp^ΔEx8/ΔEx8^* mice. *mean ± standard error of the mean. n is the number of animals.

| Cell Type | Hair Cells | Supporting Cells |
| --- | --- | --- |
| *Triobp*<sup>+/+</sup> | -29±20<br>(n=4) | 17±6<br>(n=4) |
| *Triobp*<sup>ΔEx9-10/+</sup> | -7±22<br>(n=5) | 10±1<br>(n=5) |
| *Triobp*<sup>ΔEx8/+</sup> | -10±11<br>(n=5) | 7±0.2<br>(n=5) |
| *Triobp*<sup>ΔEx9-10/ΔEx9-10</sup> | -9±10<br>(n=5) | 5±0.5<br>(n=5) |
| *Triobp*<sup>ΔEx8/ΔEx8</sup> | 4±14<br>(n=5) | 4±1<br>(n=5) |

\*mean ± standard error of the mean. n is the number of animals.

## Notes

### Competing Interest Statement

The authors have declared no competing interest.

